# Mating strategies in *Caenorhabditis elegans* populations are determined by male developmental history

**DOI:** 10.1101/2022.09.23.509199

**Authors:** Rose S. Al-Saadi, Jintao Luo, Alexandra M. Nichitean, Nikolaus R. Wagner, Douglas S. Portman, Sarah E. Hall

**Affiliations:** Department of Biology, Syracuse University, Syracuse, NY; Department of Biomedical Genetics, University of Rochester, Rochester, NY

**Keywords:** *C. elegans*, pheromone, RNAi, TGF-β, mating, postdauer

## Abstract

Mating strategies, whether sexual or asexual, confer unique costs and benefits to populations and species that facilitate evolutionary processes. In wild isolates of *Caenorhabditis elegans*, mating strategies are dependent on developmental history. Outcrossing levels significantly increase when one or both parents have transiently passed through the stress-resistant dauer diapause stage. However, the molecular mechanisms of how life history alters mating strategies have not been systematically explored. Sex-specific responses to pheromones are a major driver of mating behaviors in *C. elegans*. We demonstrated previously that postdauer hermaphrodites exhibit a decreased avoidance of the pheromone ascr#3 due to the downregulation of the *osm-9* TRPV channel gene in postdauer ADL neurons. Thus, we hypothesized that altered responses to pheromones in postdauer animals could contribute to increased outcrossing. We conducted mating assays using wild type N2 Bristol, as well as *daf-3/*co*-*SMAD and *mut-16/*Mutator strains that fail to downregulate *osm-9* in postdauer hermaphrodite ADL neurons. First, we show that the outcrossing level of N2 Bristol correlated with the developmental history of males, and that postdauer males exhibited an increased ability to detect mates via pheromones compared to continuously developed males. In addition, DAF-3 plays a critical role in postdauer males to regulate mating, while playing a more minor role in hermaphrodites. Furthermore, the *mut-16* strain exhibited negligible outcrossing, and attempts to rescue the outcrossing phenotype resulted in transgenerational sterility due to germline defects. Together, our results suggest a model whereby mating strategy is driven by developmental history under combinatorial control of TGF-β and RNAi pathways.

## INTRODUCTION

Mating strategies, whether sexual or asexual, confer unique costs and benefits to populations and species that facilitate evolutionary processes. The theoretical cost of sex and its disadvantages as a mode of reproduction are far higher than that of asexual reproduction (Otto 2009). Indeed, sexual reproduction has a two-fold cost for the existence of males due to having to share half of the offspring’s genome with a mate (Maynard Smith 1971, 1978; Butlin 2002; Gibson *et al*. 2017). In addition, sexual reproduction deprives individuals of reproductive assurance and increases the risk of predation (Meirmans *et al*. 2012; Lehtonen *et al*. 2012). Despite these disadvantages, most reproducing organisms utilize sexual reproduction to some degree. The main advantage that allowed sex to evolve and persist as the major mode of reproduction is meiotic recombination, which allows for the generation of allelic diversity, purge of deleterious mutations, and fixation of beneficial genetic variants (Weismann 1889; Burt 2000; Whitlock 2000; Agrawal 2001; Siller 2001; Paland and Lynch 2006; Whitlock and Agrawal 2009). Whether sexual reproduction is more favorable than self-fertilization reproduction depends largely on the interplay between the benefits and the costs of different reproductive modes in a particular environment.

While some species evolved to use one mode of reproduction, mixed-mating strategies are utilized by other species to benefit from both self-fertilization and sexual reproduction. For instance, daphnids can switch from asexual to sexual reproduction in response to environmental stresses such as low temperature or starvation (LeBlanc and Medlock 2015). Similarly, pea aphids produce wingless females during favorable conditions, then switch to producing winged, sexual morphs in conditions of low food and high population density (Müller *et al*. 2001). In addition, some plants can regulate their rate of self-fertilization based on environmental conditions including temperature, light, nutrient, and water availability (Johnson 1971; Campbell and Linskens 1984; Oakley *et al*. 2007; Levin 2012). Furthermore, certain *Rhabditis* species are trioecious and can reproduce via mating between males and females in favorable conditions, but develop as self-fertilizing hermaphrodites in unfavorable conditions (Félix 2004; Chaudhuri *et al*. 2011). However, the molecular mechanisms underlying these changes in mating strategies based on environmental conditions are not well understood.

*Caenorhabditis elegans* nematodes also exhibit changes in their mating strategies based on environmental experience. In conditions favorable for growth, *C. elegans* will develop continuously through four larval stages before becoming reproductive adults (control adults or CON). However, if *C. elegans* experience unfavorable conditions such as starvation, high temperature, or crowding after hatching, they can enter the stress-resistant larval diapause state called dauer (Cassada and Russell 1975). Once conditions improve, dauer larvae resume development to become reproductive adults (postdauers or PD) (Cassada and Russell 1975; Golden and Riddle 1982). Control and postdauer adults exhibit different transcriptional profiles that result in altered behaviors and life history traits (Hall *et al*. 2010; Sims *et al*. 2016; Ow *et al*. 2018; Webster *et al*. 2018; Bhattacharya *et al*. 2019). The sex of a *C. elegans* animal is determined by the number of X-chromosomes in the embryo, such that two encodes for a hermaphrodite and one for a male. Thus, males can occur spontaneously as a result of meiotic nondisjunction events involving the X-chromosome, or through cross-fertilization of an oocyte by X-null sperm (Hodgkin *et al*. 1979). Interestingly, passage through dauer over successive generations in wild isolates of *C. elegans*, CB4856 and JU440, significantly increased the proportion of males in the population and the level of outcrossing between hermaphrodites and males (Morran *et al*. 2009). Thus, the gene expression changes that occur after passage through dauer promote outcrossing behaviors in a typically self-fertilizing species.

*C. elegans* animals secrete an array of pheromone molecules which can trigger a variety of behaviors and developmental decisions such as dauer entry, aggregation, and mating (Ludewig and Schroeder 2013). Hermaphrodites produce a blend of ascaroside (ascr) molecules, including ascr#2, ascr#3, and ascr#4, which act synergistically to make up the pheromone “mating signal” at low concentrations (Srinivasan *et al*. 2008). These pheromones can elicit sexually dimorphic responses such that hermaphrodites and males exhibit avoidance or attraction behaviors, respectively, in response to high concentrations of ascr#3 (Srinivasan *et al*. 2008; Jang *et al*. 2012). Hermaphrodites acutely avoid ascr#3 via the ADL sensory neurons that connect with the backward movement neurons, AVA and AVD, via chemical synapses (Jang *et al*. 2012). In contrast, males possess a modified hermaphrodite circuit that suppresses the ADL-mediated avoidance and promotes ascr#3 attraction through ASK, ADF, and the male-specific CEM neurons (Srinivasan *et al*. 2008; Jang et al. 2012; Fagan *et al*. 2018).

These sex-specific behavioral responses depend on the expression of the OSM-9 and OCR-2 TRPV channels, which act in the signal transduction pathway downstream of pheromone receptors in amphid sensory neurons (Bargmann 2006; Jang *et al*. 2012). The expression of the *osm-9* TRPV gene in ADL neurons of hermaphrodites is dependent upon developmental history (Sims *et al*. 2016). In postdauer hermaphrodites, *osm-9* is significantly downregulated in ADL neurons, resulting in decreased avoidance in response to high concentrations of ascr#3. The altered expression of *osm-9* in postdauer hermaphrodites is regulated transcriptionally through a novel regulatory sequence called the “PD motif”. We demonstrated previously that the DAF-3 co-SMAD in the TGF-β signaling pathway and the ZFP-1/AF10 chromatin binding protein bind to the *osm-9* PD motif in postdauer ADL neurons to promote its downregulation. Interestingly, RNAi pathways are required for DAF-3 and ZFP-1 to bind at the *osm-9* promoter, suggesting that RNAi may trigger its downregulation after dauer (Sims *et al*. 2016).

Given that mating is facilitated by pheromone detection, we tested the hypothesis that sex-specific altered responses to pheromones due to altered *osm-9* expression promotes changes in mating strategies in *C. elegans* postdauer populations. Here, we show that the developmental history of males is the primary driver of mating levels, with a minor role for the developmental history of hermaphrodites. We found that postdauer males do not have altered expression of *osm-9* in ADL neurons, but do exhibit an increased ability to locate mates via pheromones compared to control males. In addition, DAF-3/co-SMAD is required in both males and hermaphrodites to regulate outcrossing levels following dauer. Similarly, we found that RNAi protein MUT-16 is also required for increased mating in postdauer populations. Attempts to rescue MUT-16 function in neurons uncovered a mechanism that promotes germline defects and transgenerational sterility. Together, our results show that *C. elegans* animals engage multiple mechanisms to increase the outcrossing frequency in postdauer populations.

## MATERIALS AND METHODS

### Strain maintenance

The strains used in this study were grown on standard NGM plates seeded with *Escherichia coli* OP50 and cultivated at 20°C unless specified otherwise (Table S1). *mut-16(pk710)* and neuronal overexpression strains were maintained at 15°C due to the temperature-sensitive mortal germline phenotype.

Control animals were maintained on plates where they had plentiful food and low population density for at least three generations. Dauers were induced by growing an age-mixed population on 60mm NGM plates until food depletion. Approximately 3-5 days later, plates were visually examined for dauers, then washed with 1% SDS to isolate the dauer larvae (Karp 2016). *daf-3(mgDf90)* exhibits a dauer-deficient phenotype and was grown on 100mm NGM plates seeded with concentrated bacteria (20x OP50) at 25°C for a week to induce dauer formation. All dauer larvae were rescued on NGM plates with food and maintained at 20°C.

### *mut-16* overexpression strains

The *mut-16* expression transgene driven by the *rab-3*, *sre-1*, *gpa-4*, and *srh-142* promoters were maintained as extrachromosomal arrays by selecting for animals that exhibit the *unc-122*p::*dsRed* (*array+*) coinjection marker as described in Bharadwaj and Hall (2017). Due to low transmission rates of the extrachromosomal transgene, we integrated the transgene array into the genome following a UV-irradiation protocol for the pan-neuronal (*rab-3*p) and ASI (*gpa-4*p) *mut-16* overexpression strains (Evans 2006). The SEH378 strain was made by injecting germ lines of N2 Bristol hermaphrodites with the *rab-3*p::*mut-16*::*gfp* and *unc-122*p::*dsRed* plasmids together at 8 ng/µL and 30 ng/µL, respectively, and selecting for *array+* progeny.

### Mating assays

Mating assays were conducted in the following combinations of continuously developed control (CON) and postdauer (PD) adult males (M) and hermaphrodites (H): CON-H x CON-M, CON-H x PD-M, PD-H x CON-M, and PD-H x PD-M. For most assays, six L4 stage hermaphrodites and six young adult males were allowed to mate on a single plate (Table S2). The parents were placed on 60mm NGM plates and allowed to mate for 23 hours, followed by removal of males from the plates. Hermaphrodites were then placed on new plates daily until egg-laying ceased and the number of progeny (hermaphrodites and males) per assay was determined. At least three mating assays using biologically independent populations were performed for each condition.

The mating frequency was determined using the proportion of males in the F1 population after mating. Since males are produced through either mating or nondisjunction events of the X-chromosome, the mating frequency can be determined by subtracting the frequency of X-chromosome nondisjunction in that strain from the overall observed male frequency (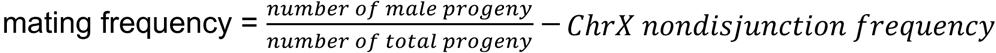). To determine the X-chromosome nondisjunction frequencies, we conducted brood size assays using at least three biological replicates of each genotype, with 6 hermaphrodites tested for each trial. The proportion of males in the progeny of each trial was averaged across replicates (Figure S1; Table S3). There was no significant difference between CON and PD male frequencies; thus, CON male frequency values were subtracted from either condition. To compare the mating assay results of the different conditions, a two-way ANOVA followed by Tukey’s multiple comparisons test was performed using GraphPad Prism version 8.4.3 for macOS, GraphPad Software, San Diego, California USA.

### Measuring *osm-9* expression levels

The expression levels of *osm-9* were measured using an integrated transgene array expressing GFP under the regulation of 350 bp of *osm-9* upstream regulatory sequences (*pdrIs1*) as previously described (Sims *et al*. 2016). The expression of GFP was determined in control and postdauer adults for wild-type and *mut-16(mg461)* strains 24 hours after the L4 larval stage. Animals were dye-filled using DiD to identify the location of ADL and mounted on agar pads with sodium azide. GFP expression was visualized using a Leica DM5500B microscope with a Hamamatsu 741 camera controller C10600 ORCA-R2. At least 44 animals over a minimum of three biologically independent trials were examined for each condition and genotype. Statistical analysis was performed using multiple *t*-tests followed by Holm-Sidak correction within GraphPad Prism (version 8.4.3).

### Pheromone avoidance assays

One-day old adult, control and postdauer males and hermaphrodites were tested for avoidance behavior in response to 100 nM ascr#3 as previously described (Hilliard *et al*. 2002; Jang *et al*. 2012; Sims *et al*. 2016). Avoidance of 1M glycerol and M13 buffer was tested as the positive and negative controls, respectively. Briefly, animals were picked onto a 60mm NGM plate without food and allowed to acclimate for 10 minutes. The drop test acute avoidance response assay was performed using a mouth pipette with a pulled glass needle. An avoidance response was recorded when a forward-crawling animal reversed within two seconds after being exposed to the stimulus. The proportion of animals responding to ascr#3 was normalized by subtracting the proportion of animals responding to the negative control (M13 buffer) as previously described (Sims *et al*. 2016). Statistical analysis was performed using multiple *t*-tests followed by Holm-Sidak correction within GraphPad Prism (version 8.4.3).

### Brood size assays

For the brood size assays, larval L4 hermaphrodites were placed on seeded NGM plate and transferred to new plates daily until egg-laying ceased, and the number of progeny per plate was determined. For the transgenerational brood size assay, the *mut-16* expression transgene was injected into *mut-16(pk710)* and N2 strains independently, and strains were propagated for more than ten generations before being used in experiments. A red fluorescent co-injection marker that is easily visualized was used when generating these *mut-16* overexpression strains; thus, *array+* refers to progeny that inherited the extrachromosomal *mut-16/dsRed* transgene array and *array-*refers to the animals that did not inherit the array (Bharadwaj and Hall 2017). Ten hermaphrodites carrying the extrachromosomal array overexpressing *mut-16* pan-neuronally were used to establish the parent plate (P_0_). Ten hermaphrodite progeny (either *array+* or *array-*) were then transferred to a new NGM plate to establish F1 and subsequent generations, and an additional ten individual hermaphrodites were used to calculate the average brood size. Statistical analysis was performed using multiple *t*-tests followed by Holm-Sidak correction within GraphPad Prism (version 8.4.3).

### DAPI straining

A standard DAPI staining protocol for whole animals was used to visualize germ cell nuclei (Francis *et al*. 1995; Qiao *et al*. 1995). Twenty-four hours after the L4 larval stage, animals were fixed in 100% cold methanol, washed with 0.1% PBST, and DAPI stained using a 1:1000 dilution. Animals were mounted on glass slides and imaged using a Leica DM5500B microscope with a Hamamatsu 741 camera controller C10600 ORCA-R2.

### Germline morphology scoring

The whole-mount DAPI stained images of each gonad were visually scored based on several criteria, with animals given a score of zero or one based on the absence or presence of each criterion. The different categories were as follows: distal tip cell (DTC) migration defects, egg-laying defects, low gamete number, abnormal gonad morphology, and germline defects. Animals scored for the presence of DTC migration defects include weak to severe meandering on dorsal pathfinding defects (Wong and Schwarzbauer 2012). The one-day old adult hermaphrodites were scored for potentially having an egg-laying defect if they had a greater number of eggs retained within the uterus in a disorganized manner at the time of staining compared to wild type (Soto *et al*. 2002). Animals were scored as having a gamete number defect by visually comparing the amount of sperm present between wild-type and mutant animals, which included low to no sperm present. Within the abnormal gonad morphology category, animals were marked for having a smaller overall gonad size and a thinner appearance of the distal gonad compared to wild-type nematodes. Germline defects were quantified by counting the total germ cell nuclei for each gonad arm using the multi-point tool in ImageJ (NIH). The germ cell nuclei were further classified into mitotic, transition, and pachytene zones by identifying the characteristic germ cell morphology. The transition and pachytene zones were deemed to start when at least two cells in a row showed the crescent-shape and basket-shape nuclei morphology, respectively (Shakes *et al*. 2009). Three trials of biologically independent populations were examined. Statistical analysis was performed using a one-way ANOVA followed by Tukey’s multiple comparisons test within GraphPad Prism (version 8.4.3).

### Quadrant mate preference and chemotaxis assays

Mate preference assays were conducted in a *him-5* background to provide sufficient males for testing. To test whether males have heightened attraction to hermaphrodites following dauer diapause, CON and PD males were used as the tester animals and immobilized hermaphrodites were used as targets. Each quadrant contained four target animals, with each quadrant alternating between the *unc-54; him-5* and *unc-54; daf-22; him-5* strains. Ten tester animals were placed in the center of the plate and allowed to move freely about the different quadrants. The location of the tester animals was noted every 30 minutes for up to 90 minutes, and the mate preference index was then calculated as previously described (Fagan *et al*. 2018). A total of 180 tester animals over 18 trials were examined for this assay. To test if hermaphrodites had an increased attraction to postdauer males, the quadrant assay was used with CON or PD *unc-54; him-5* males alternating as the targets. CON hermaphrodites were used as the tester animals. A total of 160 animals over 16 trials were tested.

To examine ascr#3 attraction, the quadrant assay was performed using ascr#3 and ethanol diluted in water as the alternating targets, and CON and PD males as the testers. The concentrations of ascr#3 tested, 100 nM and 10 nM, were chosen to evoke a weak response in control males and not limit the responses of postdauer males due to ceiling effects. The chemotaxis index was calculated by dividing the number of animals in the ascr#3 quadrants by the total number of animals in all quadrants (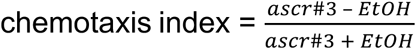). A total of 100 animals over 10 trials were tested in this assay. Statistical analysis was performed using unpaired *t*-tests within GraphPad Prism (version 8.4.3).

## RESULTS

### Outcrossing levels in N2 Bristol populations are dependent on the developmental history of males

To investigate the molecular mechanisms that promote outcrossing in postdauer populations, we first investigated whether the common *C. elegans* lab strain, N2 Bristol, exhibits an increased outcrossing frequency after passage through dauer as observed previously for other wild isolates, CB4856 and JU440 (Morran *et al*. 2009). We performed mating assays with all combinations of CON and PD, male and hermaphrodite N2 Bristol adults. Males develop from embryos that are haploid for the X chromosome, which result from meiotic chromosome nondisjunction events or fertilization of oocytes by sperm that lack the X-chromosome (Hodgkin *et al*. 1979). Thus, the outcrossing frequency was calculated using the proportion of male progeny resulting from the mating assay minus the X-chromosome nondisjunction rate as previously described (Morran *et al*. 2009; see Materials and Methods). For mating assays including CON hermaphrodites and CON males, we observed an average male frequency of 0.172 ± 0.048 among the F1 progeny, serving as the baseline of outcrossing levels (Figure 1A). When either sex experienced dauer (PD males or PD hermaphrodites), we observed a slight, albeit nonsignificant, increase in outcrossing levels (0.293 ± 0.035 and 0.209 ± 0.036, respectively) compared to the baseline (Figure 1A). However, when both sexes had experienced dauer (PD males and PD hermaphrodites), we observed a significant increase in outcrossing (0.350 ± 0.025) compared to the mating assays using CON adults (*p* = 0.012) (Figure 1A). We found that the type of males (CON or PD), but not the type of hermaphrodites, had a significant effect on outcrossing levels in N2 Bristol (*p* = 0.0016, two-way ANOVA) (Table S4), indicating that the developmental history of males is the major determinant of outcrossing in wild-type populations. However, when comparing the outcrossing levels from assays where only males experienced dauer to the assays where neither parent or both parents experienced dauer, no significant differences were observed (Figure 1A). This observation suggests a minor role for the hermaphrodite’s developmental history in promoting facultative outcrossing in populations that transiently experienced dauer diapause. Together, these results demonstrate that postdauer populations of N2 Bristol exhibit increased outcrossing levels similar to the CB4856 (Hawaiian) isolates reported previously and that the developmental history of males is the main determinant of outcrossing levels.

**Figure 1.**
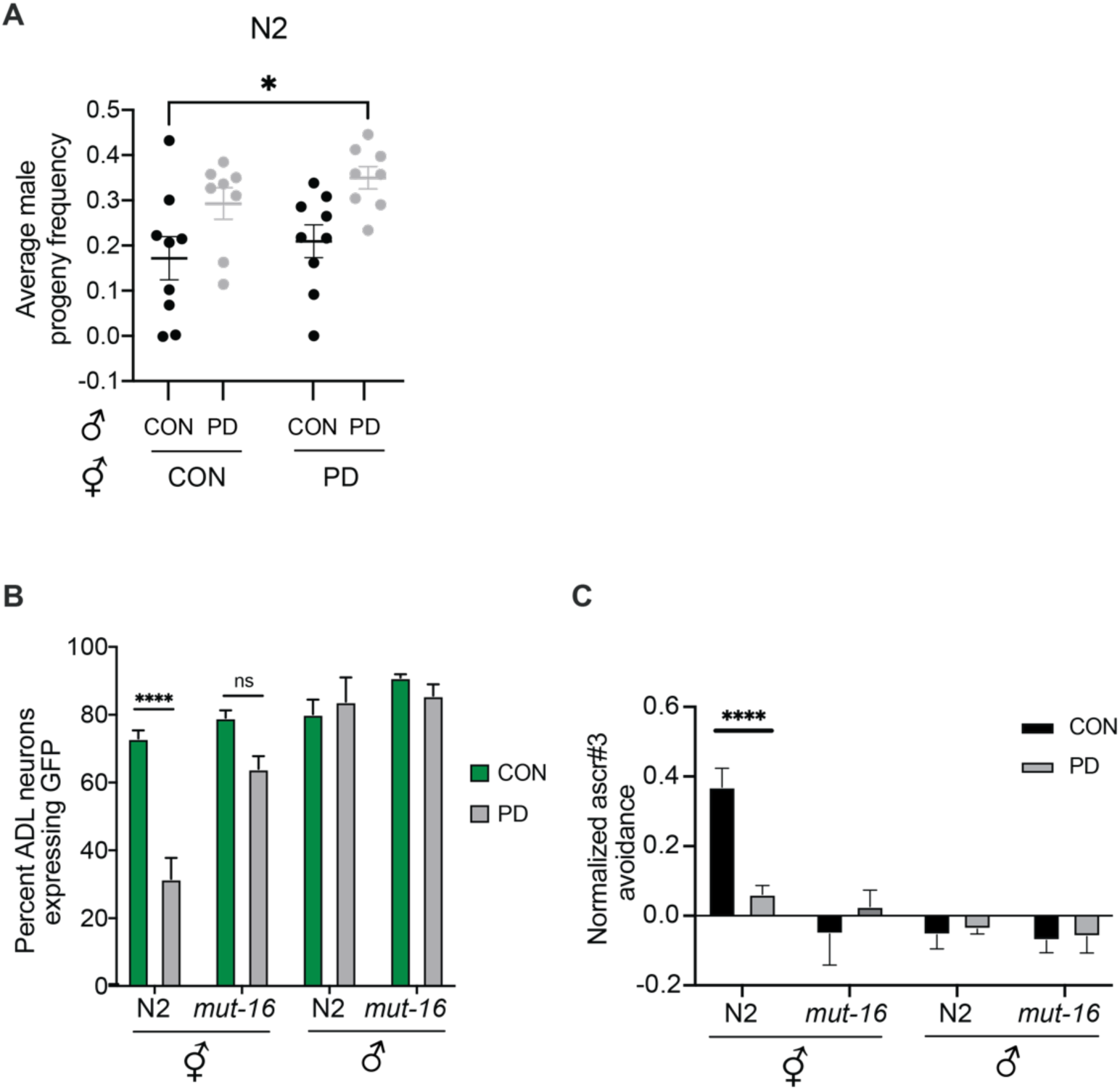
Outcrossing levels in N2 Bristol are dependent upon the developmental history of the males. **(A)** Scatter plots with lines representing mean ± S.E.M. male progeny frequencies resulting from mating assays including combinations of CON and PD adults for wild type N2. * *p* < 0.05, two-way ANOVA with Tukey’s *post hoc* test. Trial and sample sizes are provided in Table S2. Additional data are provided in Table S4. **(B)** Mean percentage ± S.E.M. of ADL neurons expressing GFP in N2 and *mut-16*(*mg461*) strains. N ≥ 44 animals from at least 3 independent trials. **** *p* < 0.0001, multiple *t*-tests followed by Holm-Sidak correction. n.s., not significant. **(C)** Normalized mean ascr#3 avoidance ± S.E.M. for hermaphrodites and males in N2 and *mut-16(pk710)* strains. n ≥ 93 animals from at least 3 independent trials. **** *p* < 0.0001, multiple *t*-tests followed by Holm-Sidak correction. Additional data are provided in Table S5.

### Expression of *osm-9* is regulated in a sex-specific manner

Since our original hypothesis was that variable *osm-9* expression altered pheromone responses resulting in increased postdauer outcrossing rates, we next examined the regulation of *osm-9* in males. Using a strain carrying an integrated GFP reporter transgene driven by ∼350 bp of the *osm-9* promoter (*osm-9*p*::gfp*), we quantified the number of ADL neurons that express GFP in control and postdauer wild-type hermaphrodites and males. As expected, we observed that significantly fewer postdauer hermaphrodites have *osm-9*p::*gfp* expression in ADL compared to controls (Figure 1B). However, in males, GFP is expressed similarly in control and postdauer adult ADL neurons, indicating that regulation of *osm-9* by developmental history is sex-specific (Figure 1B). We showed previously that hermaphrodites carrying a mutation in *mut-16* exhibited increased expression of GFP in postdauer hermaphrodite ADL neurons compared to wild type (Sims *et al*. 2016). MUT-16 is a glutamine/asparagine motif-rich protein that is required for the formation of the *Mutator* focus, a phase-separated perinuclear condensate important for small interfering RNA (siRNA) amplification (Zhang *et al*. 2011; Phillips *et al*. 2012; Uebel *et al*. 2018). To examine if MUT-16 also plays a role in the regulation of *osm-9* expression in males, we examined the GFP expression in ADL neurons in control and postdauer males in the *mut-16* strains. We observed that GFP continued to be expressed in control and postdauer males in the *mut-16* strain similar to wild type (Figure 1B), indicating that MUT-16 does not play a role in regulating *osm-9* expression in male ADL neurons.

Avoidance of ascr#3 is an ADL-mediated, *osm-9* dependent behavior in hermaphrodites (Jang *et al*. 2012). We demonstrated previously that postdauer hermaphrodites fail to avoid high concentrations of ascr#3 due to the downregulation of *osm-9* in their ADL neurons (Sims *et al*. 2016). To examine the ascr#3 avoidance behavior in postdauer males, we tested the response of wild-type and *mut-16* control and postdauer hermaphrodites and males to 100 nM ascr#3 using a drop test assay, as well as M13 buffer (negative control) and 1M glycerol (positive control) (Figure 1C; Figure S2). While the positive and negative controls elicited the expected behaviors in both control and postdauer males for the N2 and *mut-16(pk710)* strains (Figure S2), exposure to ascr#3 did not elicit a difference in avoidance behavior in postdauer males compared to control males (Figure 1C). Together, these results indicate that the differential regulation of *osm-9* in ADL neurons due to passage through the dauer stage is hermaphrodite-specific and is unlikely to be a significant driver of outcrossing in postdauer males.

### Endogenous RNAi regulates mating frequency

Since the endogenous RNAi pathway is required for the downregulation of *osm-9* in postdauer hermaphrodite ADL neurons (Sims *et al*. 2016), we tested the hypothesis that RNAi is also required for increased outcrossing in postdauer populations by performing mating assays using the *mut-16(pk710)* mutant strain. In the mating assay of *mut-16* control hermaphrodites and control males, we found an average male progeny frequency similar to what was observed for wild type (0.129 ± 0.028) (Figure 2A). However, unlike wild type, we found that the male progeny frequencies from all mating assay combinations of *mut-16* control and postdauer adults were not significantly different and were similar to the baseline outcrossing levels (Figure 2A). These results indicate that MUT-16 is required for the increased outcrossing phenotype in postdauer populations.

**Figure 2.**
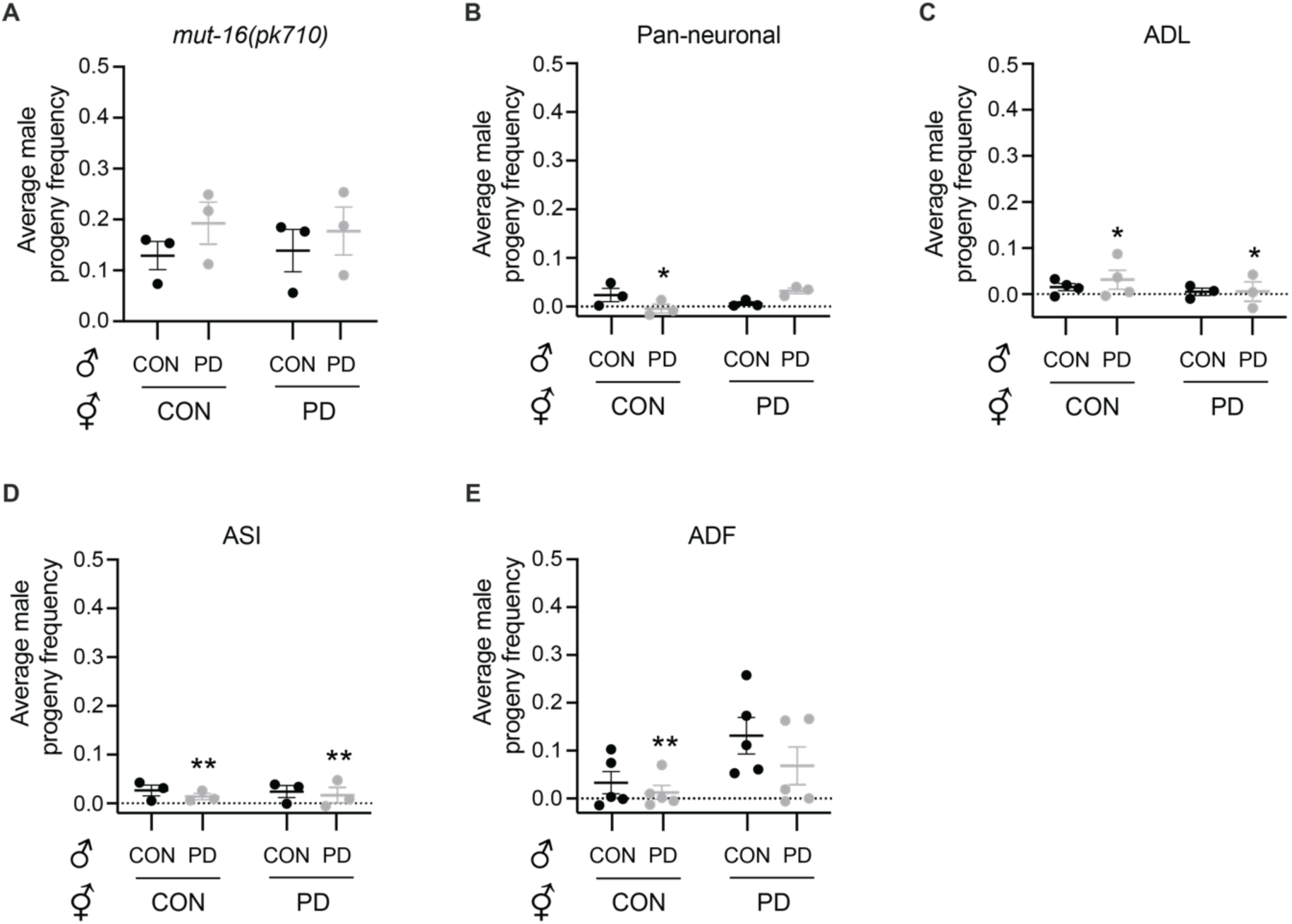
Endogenous RNAi is required for increased outcrossing in postdauer populations. Scatterplots with lines representing mean ± S.E.M. male progeny frequencies resulting from mating assays including combinations of CON and PD adults for **(A)** *mut-16(pk710)* alone or *mut-16(pk710)* animals carrying a *mut-16* transgene expressed **(B)** pan-neuronally (*rab-3*p), **(C)** in ADL neurons (*sre-1*p), **(D)** in ASI neurons (*gpa-4*p), and **(E)** in ADF neurons (*srh-142*p). Mating assays within a strain were not significantly different. * *p* < 0.05, ** *p* < 0.01, comparison of same mating assay to *mut-16(pk710)* using two-way ANOVA followed by Tukey’s multiple comparisons test. Trial and sample sizes are provided in Table S2. Additional data are provided in Tables S6-S10.

In previous work, we demonstrated that *mut-16* animals were partially dauer deficient in the presence of high pheromone levels due to decreased expression of genes encoding G proteins required for neuronal signaling. In addition, we were able to rescue the dauer deficient larval phenotype by expressing a high-copy transgene array carrying wild-type *mut-16* coding sequence either pan-neuronally or in individual, pheromone-sensing neurons (Bharadwaj and Hall 2017). To determine if MUT-16 was also functioning in neurons to regulate mating behaviors, we performed the mating assay using a *mut-16* strain carrying a *mut-16* expression transgene driven by a pan-neuronal promoter (*rab-3*p). Interestingly, we found the frequencies of male progeny in the *mut-16* pan-neuronal overexpression strain mating assays were suppressed below baseline outcrossing in *mut-16* mutants, particularly for the mating assay including CON hermaphrodites with PD males (Figures 2A and 2B). Next, we attempted to perform mating assays using *mut-16* strains that carry overexpression transgene arrays driven by single-neuron promoters. We examined the male progeny frequencies of *mut-16* strains carrying the *mut-16* transgene expressed only in ADL neurons (*sre-1*p) and found that this strain also exhibited suppressed outcrossing below baseline, especially for mating assays including PD males (Figure 2C). We observed similar suppressed outcrossing levels for the *mut-16* strains expressing the transgene in the pheromone-sensing ASI neurons (*gpa-4*p) (Figure 2D). However, *mut-16* overexpression in ADF neurons (*srh-142*p) did not result in as severe of mating suppression as the other neurons tested in assays that included postdauer hermaphrodites (Figure 2E). These results support the conclusion that *mut-16* expression is required at endogenous levels for promoting outcrossing in postdauer populations

### Neuronal overexpression of *mut-16* negatively influences hermaphrodite fecundity

While performing the outcrossing assays with the *mut-16* pan-neuronal and individual neuron overexpression strains, we observed that they exhibited a decrease in total progeny compared to wild-type and *mut-16* strains. To investigate if their suppressed outcrossing levels were due to germline defects, we first examined the ability of the *mut-16* neuronal overexpression strain hermaphrodites to produce self-fertilized progeny. We conducted brood size assays of CON and PD hermaphrodite adults with N2 Bristol, *mut-16*(*pk710*), and two independent lines of the *mut-16* strain carrying an extrachromosomal array of pan-neuronal *mut-16* expression transgene. First, we observed that wild-type postdauer hermaphrodites showed a significantly reduced brood size compared to controls, and that this difference is eliminated in the *mut-16(pk710)* strain as we reported previously (Figure 3A) (Hall *et al*. 2010; Ow *et al*. 2018). Next, we found that control and postdauer adults of two independent pan-neuronal *mut-16* overexpression lines exhibited a significant reduction in brood size compared to wild-type and *mut-16* strains (Figure 3A), indicating that the low fecundity of the overexpression strains was not merely due to the *pk710* allele alone. Thus, overexpression of *mut-16* in neurons has negative consequences on hermaphrodite fecundity in *mut-16* mutants.

**Figure 3.**
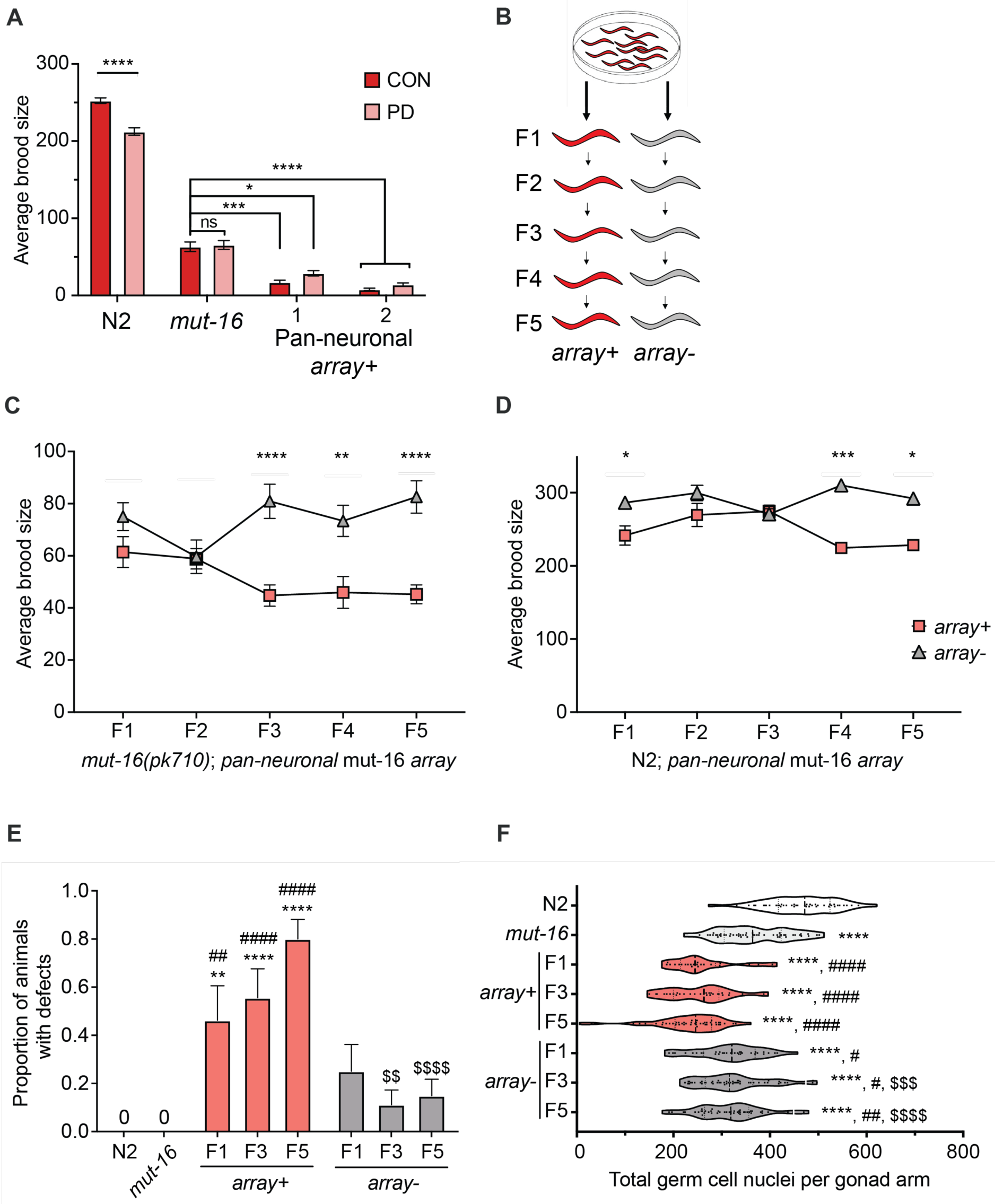
Neuronal overexpression of *mut-16* decreases hermaphrodite fecundity transgenerationally. **(A)** Bar graph represents mean brood size ± S.E.M. for CON and PD hermaphrodites of wild type N2 and *mut-16*(*pk710*) strains, and *mut-16*(*pk710*) adults carrying the pan-neuronal *mut-16* expression array. * *p* < 0.05, *** *p* < 0.001, **** *p* < 0.0001, one-way ANOVA with Tukey’s *post hoc* test. n.s. not significant. **(B)** Diagram illustrating the experimental design of transgenerational brood size assays. **(C, D)** Mean brood size ± S.E.M. for **(C)** *mut-16(pk710)* and **(D)** N2 strains carrying the pan-neuronal *mut-16* overexpression array (*array+*) or that have lost the array (*array-*) from F1 to F5 generations. * *p* < 0.05, ** *p* < 0.01, *** *p* < 0.001, **** *p* < 0.0001, multiple *t*-tests followed by Holm-Sidak correction. **(E)** Mean proportion ± S.E.M. of hermaphrodite adults with small gonads in N2, *mut-16(pk710),* and *mut-16(pk710) array+* or *array-* strains. **(F)** Violin plots represent the total number of germ cell nuclei per gonad arm for N2, *mut-16(pk710),* and *mut-16(pk710) array+* or *array-* strains. **(E, F)**, *, #, $ *p* < 0.05; **, ##, $$ *p* < 0.01; ***, ###, $$$ *p* < 0.001; ****, ####, $$$$ *p* < 0.0001; one-way ANOVA with Tukey’s *post hoc* test. * denotes a comparison to N2; # denotes a comparison to *mut-16(pk710),* $ denotes a comparison to *array+* within the same generation. Trial and sample sizes are provided in Table S2. Additional data are provided in Tables S11-S13.

### The negative effects of MUT-16 overexpression on hermaphrodite fecundity are transgenerationally inherited

Double-stranded RNA (dsRNA), the trigger for RNAi, has been shown to regulate germline gene expression transgenerationally when expressed in neurons (Devanapally *et al*. 2015). Thus, we questioned whether the reduced brood size observed in the *mut-16* pan-neuronal overexpression strains could be inherited over generations that have lost the *mut-16* expression transgene. To address this question, we counted the brood sizes of F1 to F5 generations of *mut-16* hermaphrodites that have retained (*array+*) or lost (*array-*) the extrachromosomal *mut-16* transgene array in the F1 generation from an *array+* hermaphrodite parent (see Materials and Methods) (Figure 3B). The F1 and F2 generations did not exhibit a significant difference in brood size between *array+* and *array-* populations. However, for the F3 through F5 generations, we observed that the *array-* populations had significantly larger brood sizes compared to their *array+* counterparts (Figure 3C). This result was due to a gradual increase and decrease in fecundity of the *array-* and *array+* progeny, respectively, over the five generations examined.

We next asked if the decline in hermaphrodite fecundity of the *mut-16* pan-neuronal overexpression strains was due to accumulated mutations in a sensitized *mut-16*(*pk710*) background. In *mut-16* animals, transposon activity is derepressed, resulting in increased mutagenesis and decreased genome integrity (Ketting *et al*. 1999; Tabara *et al*. 1999; Sijen and Plasterk 2003). Hypothetically, expression of MUT-16 in neurons of a *mut-16(pk710)* animal would not rescue the transposon suppression defects in other tissues, including the germ line. To address this question, we injected the wild-type N2 strain with the pan-neuronal *mut-16* expression transgene array and examined the brood sizes of *array+* and *array-* progeny of the F1 to F5 generations from an *array+* hermaphrodite. For the F1 through F3 generations, the *array+* populations show variable, but similar, brood sizes to the *array-* progeny (Figure 3D). In the F4 and F5 generations, we observed a similar separation in brood sizes compared to the *mut-16(pk710)* strain, where brood sizes of *array+* adults were significantly larger than *array-* adults (Figure 3D). Together, these results suggest that the neuronal overexpression of *mut-16* is correlated with increased sterility over generations, possibly through inheritance of epigenetic factors such as small RNAs.

### Neuronal overexpression of *mut-16* negatively influences gonad morphology and germ cell number transgenerationally

After observing the decrease in fecundity caused by overexpression of *mut-16* pan-neuronally, we investigated possible mechanisms leading to this phenotype that contributed to the appearance of decreased mating. Recently, *mut-16* mutants were shown to exhibit germline defects following heat stress, including small or collapsed gonad arms, defects in maintaining germ cell pluripotency, and misregulation of spermatogenic, oogenic, and somatic genes in the germ line (Rogers and Phillips 2020). Thus, we asked whether neuronal overexpression of *mut-16* worsened the gonad morphology defects of *mut-16* animals, leading to sterility. Since the decreased fecundity is more severe in animals retaining the array compared to *mut-16(pk710)* animals (Figure 3A), we expected to observe a correlation of gonad morphology defects with *array+* animals that were not present in the *array-* counterparts. To test this hypothesis, we examined the somatic gonad and germline morphology for defects in the F1, F3, and F5 generations of *mut-16* pan-neuronal overexpression strain hermaphrodites that either retained (*array+*) or lost (*array-*) the extrachromosomal array. As reported previously, the proportion of *mut-16(pk710)* animals displaying overall gonad defects was significantly higher than that of N2 Bristol, including DTC migration defects, decreased sperm number, and thin distal gonads (Figures S3 and S4). However, for small gonad defects, we detected a trend in the proportion of animals exhibiting this defect and the observed brood sizes of these populations. Interestingly, the *array+* populations showed an increasing proportion of defects between the F1 and F5 generations, while the *array-* populations exhibited significantly fewer defects (Figure 3E). Moreover, neither wild-type N2 nor *mut-16* strains exhibited the small gonad phenotype, suggesting that the defect was a consequence of the pan-neuronal *mut-16* array.

To further characterize how the neuronal overexpression of *mut-16* contributes to decreased outcrossing, we examined whether small gonad defects were correlated with fewer germ cells available for reproduction. Using DAPI-stained whole hermaphrodites, we quantified the number of germ cells per gonad arm of *array+* and *array-* progeny for F1, F3, and F5 generations and compared it to wild-type and *mut-16(pk710)* strains. As expected, we found that *mut-16(pk710)* hermaphrodites exhibited a significant decrease in germ cell counts compared to wild type (Figure 3F). Furthermore, we observed that *array+* generations exhibited a further decrease in total germ cells compared to *mut-16* adults, and the *array-* animals exhibited an intermediate germ cell number between the *mut-16* and *array+* populations (Figure 3F). Upon further scrutiny, we found that the differences in total germ cell number appear to be due to a decrease in cells in the mitotic and pachytene regions of the germ line (Figures S5 and S6). Together, these results indicate that the overexpression of MUT-16 in neurons can negatively impact hermaphrodite fecundity and germ cell numbers transgenerationally. Due to the heritable, negative effects of *mut-16* overexpression on fecundity and germline morphology, we were unable to further investigate the role of MUT-16 in outcrossing levels using these strains.

### TGF-β signaling pathway is required for increased outcrossing in postdauer populations

We previously showed that the *osm-9* TRPV channel gene is regulated by the TGF-β pathway in ADL neurons of postdauer hermaphrodites. Specifically, the DAF-3/co-SMAD transcription factor binds to the *osm-9* promoter to promote its downregulation after dauer, and mutations in *daf-3* result in continued *osm-9* expression and ascr#3 avoidance in postdauer hermaphrodites (Sims *et al*. 2016). Thus, we tested whether the increased outcrossing levels in postdauer populations were dependent upon TGF-β signaling by performing mating assays using a *daf-3(mgDf90)* mutant strain. For mating assays of *daf-3* CON hermaphrodites and CON males, we observed a similar baseline of male progeny frequency to what we observed for wild type (0.154 ± 0.040) (Figures 4A). Although we did observe a slight increase in outcrossing when the *daf-3* mating assays included postdauer males, postdauer hermaphrodites, or both, the male progeny frequencies were not significantly different from baseline (Figure 4A; Table S14). These results suggest that DAF-3 and the TGF-β signaling pathway may play a minor role in the increased outcrossing in postdauer populations. Since *osm-9* is only downregulated in postdauer hermaphrodite ADL neurons, we next asked whether DAF-3 was required in males or hermaphrodites to promote postdauer outcrossing. We predicted that the continued expression of *osm-9* and ascr#3 avoidance in postdauer *daf-3* hermaphrodites would decrease outcrossing. To test our hypothesis, we examined male frequencies in the progeny of populations where only one sex carrying a mutant allele of *daf-3* was mated with wild-type animals of the other sex. In populations where *daf-3* males were mated to wild-type hermaphrodites, we found that the trends in outcrossing were the opposite compared to wild type. Mating assays that included PD males showed a trend of decreased male progeny frequencies compared to mating assays that included CON males. Interestingly, mating assays that included PD wild-type hermaphrodites and CON *daf-3* males exhibited significantly higher male progeny frequencies compared to mating assays with CON wild-type hermaphrodites and PD *daf-3* males (*p* = 0.016; Figure 4B). These results are consistent with a requirement for DAF-3 in postdauer males to promote outcrossing.

**Figure 4.**
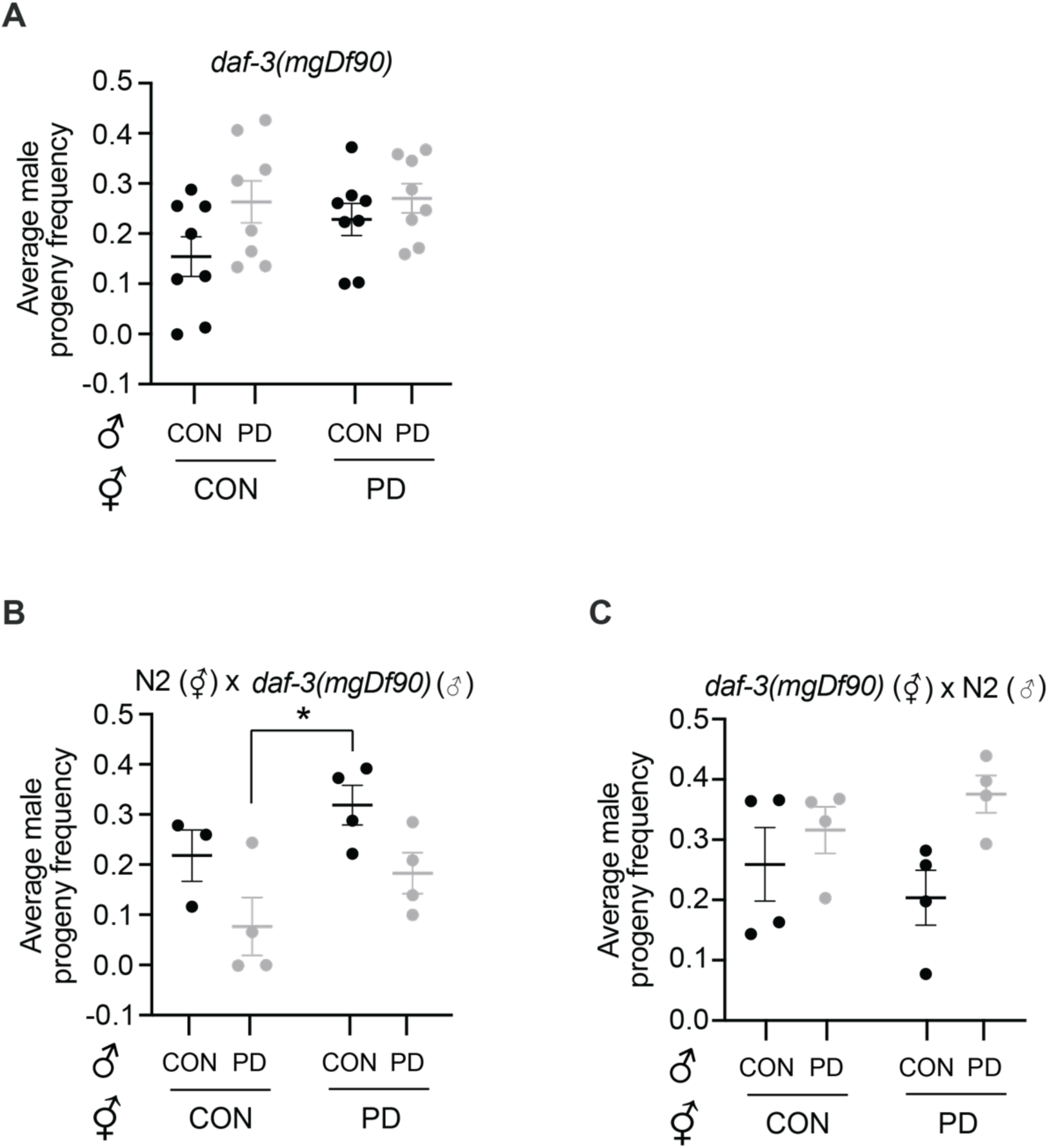
TGF-β signaling plays a role in increased outcrossing levels in postdauer populations. Scatterplots with lines representing the average male progeny frequencies ± S.E.M. for mating assays including CON and PD populations of **(A)** *daf-3*(*mgDf90*), **(B)** N2 hermaphrodites mated with *daf-3* males, and **(C)** *daf-3* hermaphrodites mated with N2 males. * *p* < 0.05, two-way ANOVA with Tukey’s *post hoc* test. Trial and sample sizes are provided in Table S2. Additional data provided in Tables S14-S16.

We further investigated a potential role of DAF-3 in hermaphrodites by examining the male progeny frequency in mating assays of *daf-3* hermaphrodites with wild-type males. If hermaphrodites did not play a significant role in outcrossing levels, we would expect that male progeny frequencies of these assays would be similar to wild type. Instead, we observed that all combinations of CON and PD populations did not show significant differences in male progeny frequencies (Figure 4C). However, the mating assays including PD wild type males trended towards increased male progeny frequencies compared to assays with CON males similar to the *daf-3* mating assays, suggesting a more minor role for DAF-3 in hermaphrodites (Figures 4A and 4C). Together, these results indicate that DAF-3 is required in both males and hermaphrodites to promote outcrossing in postdauer populations.

### Postdauer males exhibit increased ability to detect mates via pheromone

Males find mates through detection and chemotaxis towards pheromone molecules produced by hermaphrodites (Simon and Sternberg 2002; Srinivasan *et al*. 2008; Fagan *et al*. 2018). Our results indicate that the developmental history of the male is a significant contributor to outcrossing rates, with hermaphrodites playing a more minor role. We hypothesized two non-mutually exclusive mechanisms to explain how postdauer males contribute to outcrossing levels. First, postdauer males may be more efficient at finding mates than control males. A second possibility is that postdauer males may produce a novel pheromone component that functions as an attractant to hermaphrodites.

To test these possibilities, we first performed mate preference assays using genetically immobilized hermaphrodites to determine male preference and their ability to locate mates (Fagan *et al*. 2018). Each quadrant contained immobilized control hermaphrodites that can (Phe^ON^: *unc-54*) or cannot (Phe^OFF^: *unc-54; daf-22*) secrete pheromones into their environment. In each assay, either control or postdauer wild-type males were placed in the center of the plate and scored for their location in the quadrants after 30, 60, and 90 minutes (Figure 5A). Both control and postdauer males exhibited preference towards hermaphrodites that were able to secrete pheromones (Figure 5B). Interestingly, we observed that postdauer males were better able to locate the pheromone-producing hermaphrodites compared to control males (Figure 5B). These results support the hypothesis that postdauer males are more efficient at locating mates compared to control males. To test if the increased postdauer male efficiency at locating mates was facilitated by ascr#3 detection, we also conducted chemotaxis quadrant assays with 10 nM and 100 nM ascr#3 (Fagan *et al*. 2018). Males were tested for their preference to four quadrants containing either ascr#3 or ethanol (Figure 5C). Postdauer males exhibited a slight, but statistically insignificant, increase in chemotaxis towards both 10 nM and 100 nM ascr#3 concentrations compared to control males (Figure 5D). This result was not surprising given that males are attracted to a blend of pheromones secreted by hermaphrodites, and ascr#3 detection may be just one contributor to the increased mate finding efficiency of postdauer males.

**Figure 5.**
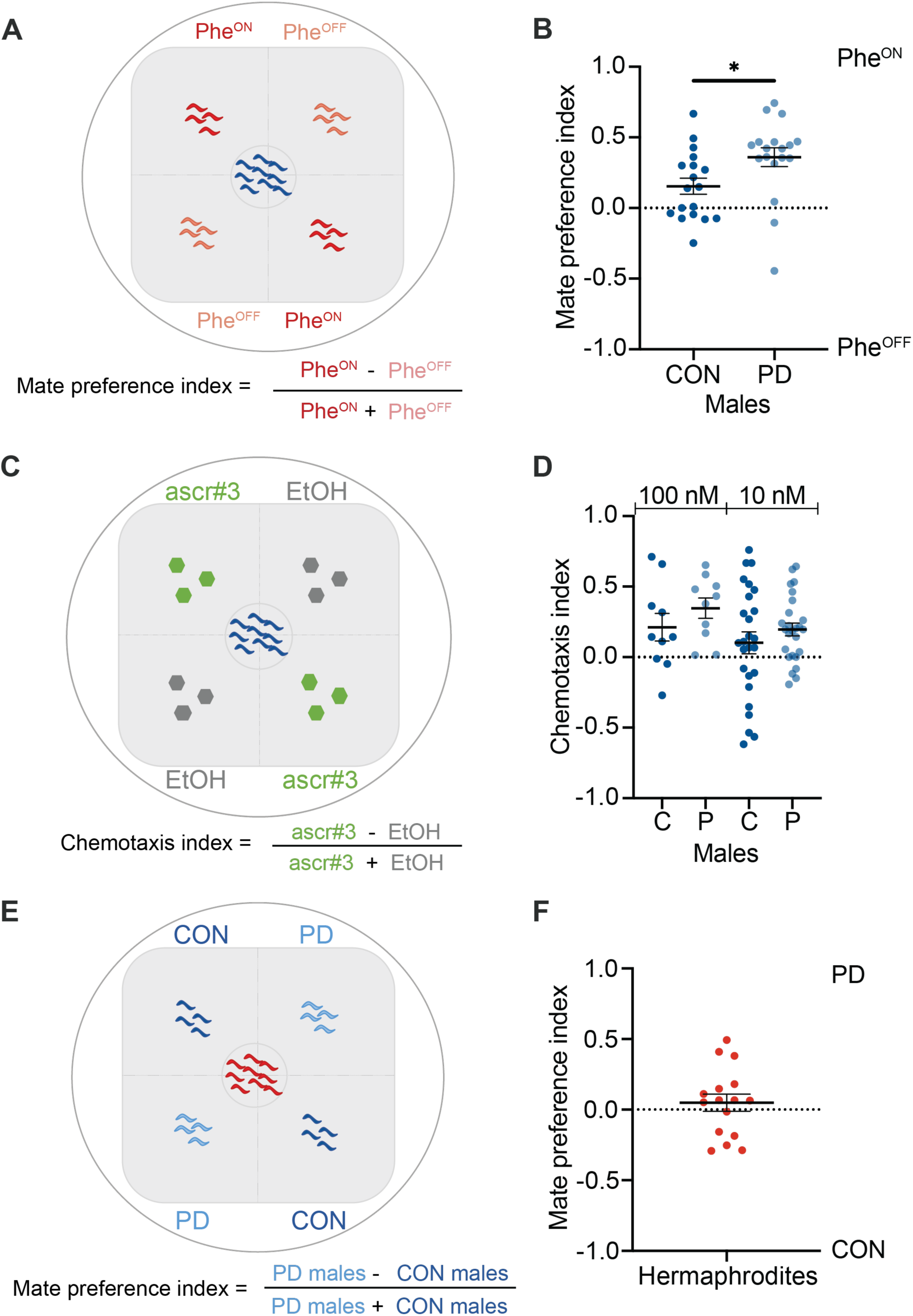
Postdauer males exhibit an increased ability to locate mates via pheromone. **(A, E)** Schematic for the mate preference and **(C)** chemotaxis assay experimental setups. **(A)** Immobilized hermaphrodites of the same genotype (Phe^ON^: *unc-54* or Phe^OFF^: *unc-54; daf-22*) were placed on opposite quadrants and, **(B)** males were scored based on quadrant location. **(C, D)** CON and PD *him-5* male preference for ascr#3 or EtOH quadrants. **(E)** Immobilized CON and PD *him-5* males were placed in opposite quadrants and **(F)** hermaphrodites were scored based on quadrant location. N ≥ 10 trials; n ≥ 100 animals. * *p* < 0.05, unpaired *t*-test. Additional data are provided in Table S17.

To test the possibility that postdauer males secrete a pheromone for hermaphrodite attraction, we again performed the mate preference quadrant assays using immobilized (*unc-54*) control or postdauer males in each quadrant. The location of control wild-type hermaphrodites was scored every 30 minutes, up to 90 minutes (Figure 5E). In contrast to the assay for male preference, hermaphrodites did not exhibit any preference between control and postdauer males (Figure 5F). Together, these results suggest that postdauer males promote outcrossing through their increased ability to locate potential mates using pheromone cues.

## DISCUSSION

Phenotypic plasticity plays a significant role in regulating reproductive modes. Here, we showed that the change in reproductive strategies to increase mating in *C. elegans* following dauer diapause requires sexually dimorphic changes in behavior. First, the developmental history of males was the major contributor to outcrossing levels, potentially due to the increased ability of postdauer males to detect mates using pheromone cues compared to control males. In addition, we uncovered a minor role for the developmental history of hermaphrodites in regulating mating levels. These phenotypes depend on RNAi and TGF-β pathways, which potentially act to regulate pheromone production and detection. Furthermore, we demonstrate that overexpression of MUT-16 pan-neuronally or in individual neurons negatively regulates hermaphrodite fecundity by decreasing germ cell number transgenerationally. Together, our results illustrate some of the genetic determinants required for increased outcrossing behaviors in postdauer populations.

### Contributions of males and hermaphrodites towards outcrossing frequency

Our original hypothesis driving this work was that downregulation of *osm-9* expression in postdauer hermaphrodite ADL neurons facilitated the opportunity for increased mating due to their failure to avoid high concentrations of ascr#3. However, our results indicated that the developmental history of males was the primary determinant of outcrossing rates, with hermaphrodite life history playing a more minor role (Figure 1A). *C. elegans* males balance their drive for reproduction and survival by alternating between foraging (food-seeking) and exploration (mate-seeking) behaviors. Male exploration is suppressed by the presence of hermaphrodites and governed by the nutritional status of the males (Lipton *et al*. 2004; Wexler et al 2020). Exploratory behavior is regulated by the ASK, ASI, and AWC sensory neurons, and a change in the activity of these neurons in postdauer males could modulate the switch between foraging and exploration behaviors more efficiently or more frequently than control males, resulting in the increased outcrossing rate we observed (Gray *et al*. 2005). This hypothesis is supported by the recent evidence showing that *C. elegans* hermaphrodites of the Hawaiian wild isolate CB4856 can permanently alter their foraging behavior based on passage through the starvation-induced dauer stage (Pradhan *et al*. 2019). While this change in foraging behavior is not seen in N2 Bristol postdauer hermaphrodites, the possibility exists that males of this strain experience a sex-specific change in their foraging behavior in response to starvation-induced dauer.

Hermaphrodites influence males to switch from foraging behavior to mate-locating behavior by secreting a mixture of “mate-finding” pheromone molecules (Simon and Sternberg 2002). Males are attracted to a particular blend of ascarosides that includes ascr#2, ascr#3, ascr#4, and ascr#8, with males being most attracted to ascr#3 (Srinivasan *et al*. 2008; Kaplan *et al*. 2011). We showed in this study that postdauer males exhibited a trend of increased attraction, albeit nonsignificant, towards ascr#3 compared to control males (Figure 5D). In males, ascr#3 attraction is mediated by the ADF, ASK, and CEM sensory neurons, as well as RMG interneuron (Srinivasan *et al*. 2008, Jang *et al*. 2012, Fagan *et al*. 2018). ADL neurons, which mediate the default behavior of ascr#3 avoidance, are antagonized by ASK and ADF neurons to promote ascr#3 attraction in males (Jang *et al*. 2012; Fagan *et al*. 2018). Plasticity in the activity or connections of these neurons may contribute to the increased ability of postdauer males to locate mates. Indeed, cilia, amphid sheath cells, and electrical synapses have all been shown to remodel in dauers to contribute to dauer-specific behaviors (Albert and Riddle 1983; Procko *et al*. 2011; Schroeder *et al*. 2013; Bhattacharya *et al*. 2019). Interestingly, the fusion of amphid sheath cells in dauers, which interact with the ASI, AWC, ADF, ASK, and ADL amphid sensory neurons, is irreversible and may contribute to behavioral plasticity of chemotaxis behaviors in postdauer males (Procko *et al*. 2011).

The minor role of postdauer hermaphrodites contributing to increased outcrossing was uncovered through reciprocal crosses with wild type and *daf-3*/co-SMAD mutants (Figure 4). In wild-type populations, the mating level of postdauer hermaphrodites and males was significantly greater than control hermaphrodites and males, but that difference was lost in crosses that included *daf-3* hermaphrodites (Figures 1A and 4). In addition, the observation that *daf-3* postdauer males correlated with decreased mating in the presence of wild type hermaphrodites, but not *daf-3* hermaphrodites, suggests that both the developmental history and genetic background of each sex may interact to regulate mating levels. However, the role of *daf-3* in regulating outcrossing sex-specifically is not clear. One possible explanation is that *daf-3* mutants exhibit significantly increased path range, as well as changes in foraging and crawling frequency, compared to wild-type adults (Yemini et al. 2013). These changes in the relative time spent in exploration versus foraging modes could alter mating frequencies in crosses containing *daf-3* mutants compared to wild type. Another potential explanation is that TGF-ß signaling may alter ascaroside biosynthesis, which is regulated by diet and metabolic state (Kaplan *et al*. 2011). Postdauer hermaphrodites secrete a greater amount of ascr#4, whereas control hermaphrodites predominantly secrete ascr#2 (Kaplan *et al*. 2011). Interestingly, males linger in regions containing ascr#3 and ascr#4 longer than regions containing ascr#3 and ascr#2, suggesting that postdauer hermaphrodites may be more attractive to males compared to controls (Srinivasan *et al*. 2008). Although a direct link between *daf-3* and ascaroside biosynthesis has not been discovered to date, TGF-ß signaling has been shown to regulate diverse processes such as dafachronic acid biosynthesis, dauer formation, sperm motility, germline proliferation, and lipid metabolism based on environmental conditions (McKnight *et al*. 2014; Thomas *et al*. 1993; Gerisch and Antebi 2004; Pekar *et al*. 2017; Hussey *et al*. 2017).

### Transgenerational regulation of germ cell number by neuronal RNAi

In previous work, we demonstrated that *mut-16* hermaphrodites exhibited a dauer deficiency phenotype in high pheromone conditions due to their decrease in the expression of G proteins which are required in their chemosensory neurons to respond to environmental cues (Bharadwaj and Hall 2017). Although only hermaphrodites were examined previously, it is probable that MUT-16 is also required in males in order to appropriately respond to ascaroside molecules. Given that mating occurs through a pheromone-based mechanism, we predict that *mut-16* males are unable to mate with hermaphrodites due to their inability to detect mates via ascarosides. While overexpressing *mut-16* neuronally rescued the larval dauer deficiency, it failed to restore wild-type levels of outcrossing in adult animals and further reduced the frequency of male progeny compared to *mut-16* mutants without the transgene array (Figure 2) (Bharadwaj and Hall 2017). We hypothesize that the failure of the transgene array to rescue outcrossing in *mut-16* animals is due to its effects in the germ line rather than in neurons. MUT-16 is expressed in somatic and germline tissues and promotes formation of phase-separated condensates, called *Mutator* complexes, in the germ line (Phillips *et al*. 2012; Uebel *et al*. 2018). *Mutator* complexes are localized perinuclearly along with P granules, which function together to amplify siRNAs (Zhang *et al*. 2011; Phillips *et al*. 2012; Uebel *et al*. 2018). Interestingly, overexpression of MUT-16 in muscle has been shown to drive formation of *Mutator* complexes in those cells, suggesting the possibility that our neuronal overexpression of *mut-16* may have similar consequences (Uebel *et al*. 2018). However, how the function of neuronal MUT-16 changes when diffuse in the cytoplasm compared to localized in *Mutator* complexes remains unclear.

A second question raised by this experiment is how MUT-16 function in neurons is regulating gonad development and germ cell number. *mut-16* hermaphrodites have brood sizes that are approximately half of what is observed for wild type, and the strains overexpressing *mut-16* in neurons exhibit even further reduced broods and additional defects in gonad size and germ cell number (Figures 3, S3, S4, S5, S6) (Rogers and Phillips 2020). Double-stranded RNA expressed in neurons has been shown to regulate gene expression in the germ line transgenerationally (Devanapally *et al*. 2015). Given the role of MUT-16 in siRNA amplification, one likely explanation of our results is that overexpression of *mut-16* leads to accumulation of siRNAs in neurons that are transported to the germ line where they inappropriately alter gene expression, resulting in increased sterility (Figures 3, S3, S4, S5, S6). The mechanism of siRNA travel between tissues and what germline genes are being targeted will be addressed in future experiments.

These findings should be viewed in the context of evolution, as these changes in reproductive strategies may be driving adaptation. The changes in outcrossing rates and differential gene expression patterns are all likely to be mechanisms promoting populations to adapt to their environment by generating genetic diversity. Here, we show a genetic mechanism underlying the switch of reproductive systems from primarily self-fertilization to a mixed mating system in response to environmental stress during early development. Males are more efficient at entering dauer diapause, have increased survival while in the dauer stage, are actively attracted to dauer-inducing pheromones, and are better able to locate mates following dauer (Ailion and Thomas 2000; Morran *et al*. 2009; Ludewig and Schroeder 2013). These characteristics make males a tool for driving genetic diversity after dauer exposure. The change in reproductive strategies has a significant evolutionary implication: by allowing increased mating after stress, increased genetic diversity is promoted by shuffling alleles in the population. Since natural populations of *C. elegans* are often found in dauer, these mechanisms to increase genetic diversity in postdauer populations are likely to have profound effects on their genome evolution as well as to enable mechanisms to become more adaptive to future stressors (Frézal and Félix 2015).

## Supporting information

Al-Saadi et al Supplemental Tables

## ACKNOWLEDGEMENTS

We would like to thank Pallavi Bharadwaj for contributing some of the brood size data, Rebecca Butcher and Frank Schroeder and for providing ascr#3 pheromone, Eleanor Maine for reading a manuscript draft, and the Hall and Maine labs for helpful comments and suggestions throughout the project. Some strains were provided by the CGC, which is funded by NIH Office of Research Infrastructure Programs (P40 OD010440).

## DATA AND REAGENTS

Raw data for this study is included as Supplemental Tables associated with this manuscript. Animal strains and other reagents will be provided upon request after publication.

---

**Figure S1.**
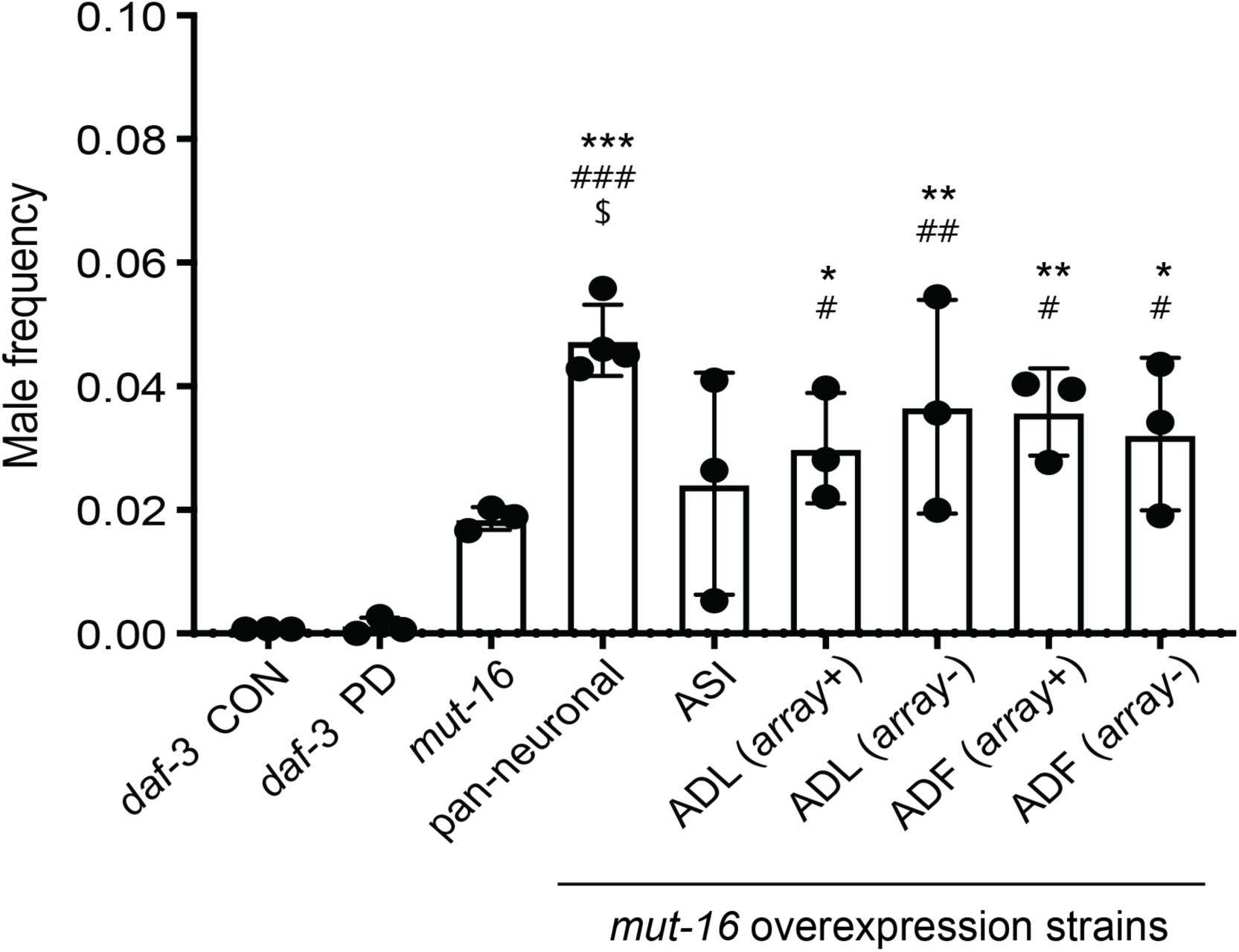
*mut-16* neuronal overexpression strains exhibit increased male incidence. Graph represents the mean male frequency ± S.E.M. n ≥ 18 animals over at least 3 biologically independent trials. The value for N2 male frequency is ≤ 0.002 as reported by (Teotónio *et al*. 2006). *, #, $ indicate *p* < 0.05; **, ## indicate *p* < 0.01; ***, ### indicate *p* < 0.001. * denotes a comparison to *daf-3* CON; # denotes a comparison *daf-3* PD; $ denotes a comparison to *mut-16*; one-way ANOVA with Tukey’s *post hoc* test. Additional data are provided in Table S3.

**Figure S2.**
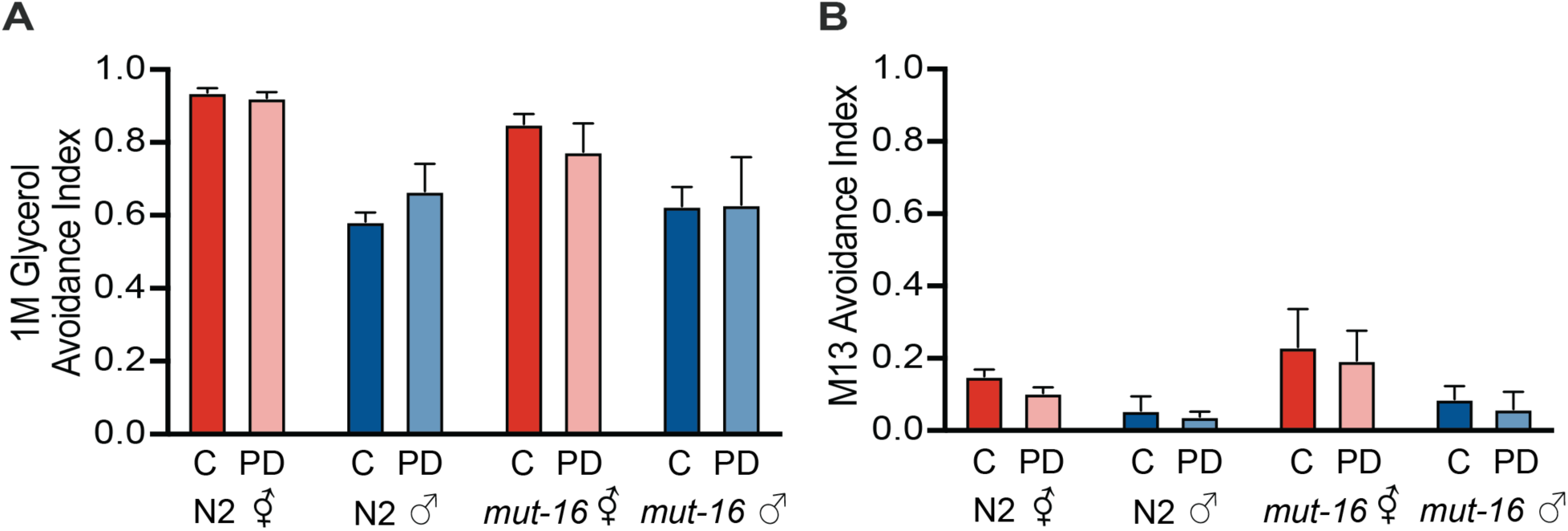
1M glycerol and M13 avoidance indices. Graphs represent mean avoidance index ± S.E.M. for wild type N2 and *mut-16(pk710)* hermaphrodites and males in response to **(A)** the positive control, 1M glycerol, and **(B)** the negative control, M13 buffer. n ≥ 76 animals over at least 3 biologically independent trials. CON and PD data within a strain are not significantly different using multiple *t*-tests. Additional data is provided in Table S5.

**Figure S3.**
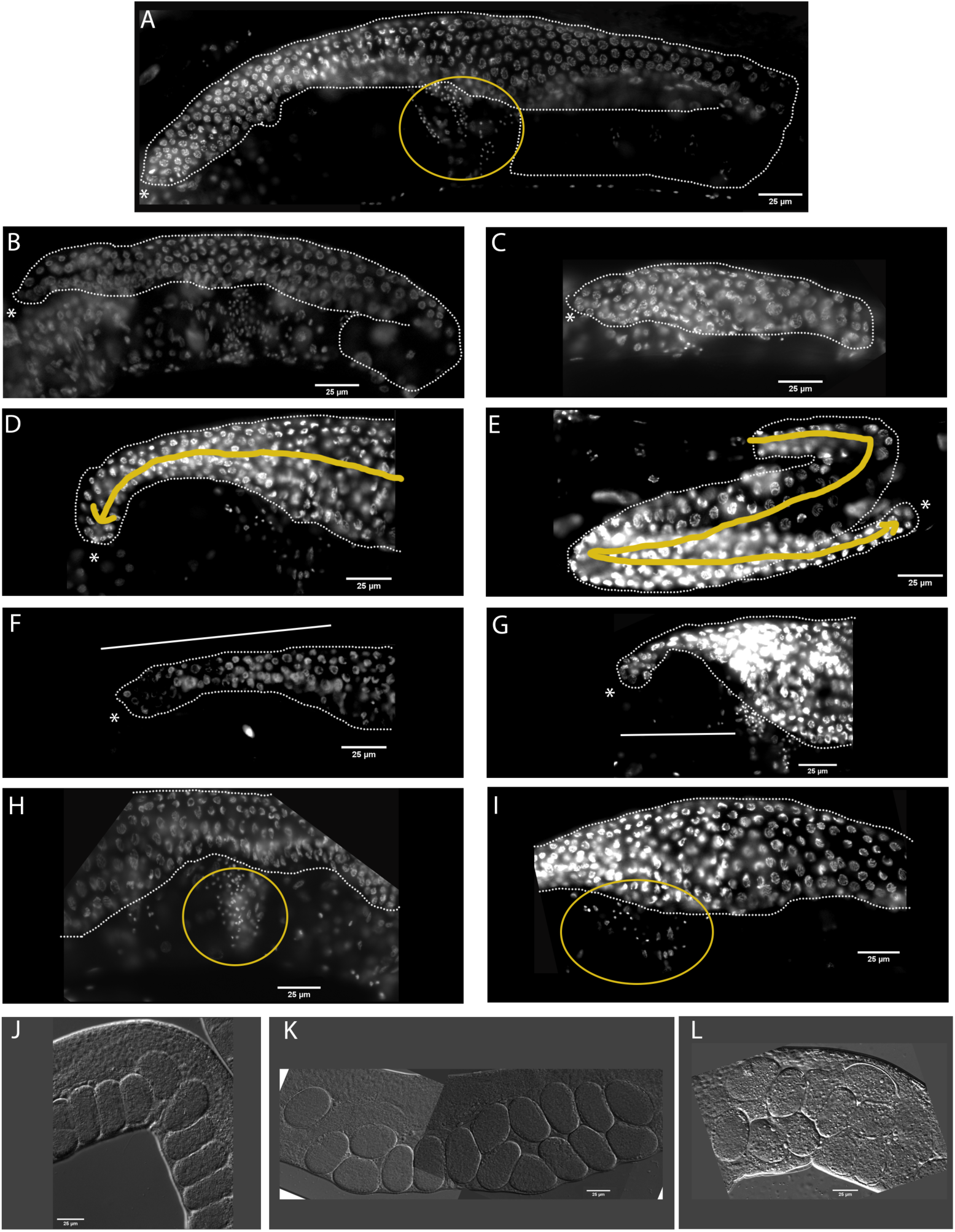
Presence of *mut-16* overexpression transgene in neurons correlates with numerous gonadal defects in *mut-16(pk710)* hermaphrodites. **(A)** Image is a representative wild-type N2 gonad. **(B, C)** Images represent **(B)** mild and **(C)** severe examples of a small gonad. **(D, E)** Images represent **(D)** mild and **(E)** severe examples of DTC migration defects. The yellow arrow indicates the probable path the DTC has taken. **(F, G)** Images represent examples of a **(F)** thin and an **(G)** extremely thin distal gonad size. The line indicates the gonad areas exhibiting the phenotype. **(H, I)** Images represent examples of **(H)** low to **(I)** extremely low sperm number. The yellow circle indicates the spermatheca. **(J)** Image represents an example of a wild-type N2 uterus. **(K, L)** Images represent examples of **(K)** mild and **(L)** severe embryo retention in *mut-16* overexpression strains. For all images, white dotted lines demark the gonad and the asterisks indicate the DTC. Scale bar is 25μm. **(A-I)** Images are oriented with the DTC towards the left side, except for **(E)** where the gonad has an additional turn.

**Figure S4.**
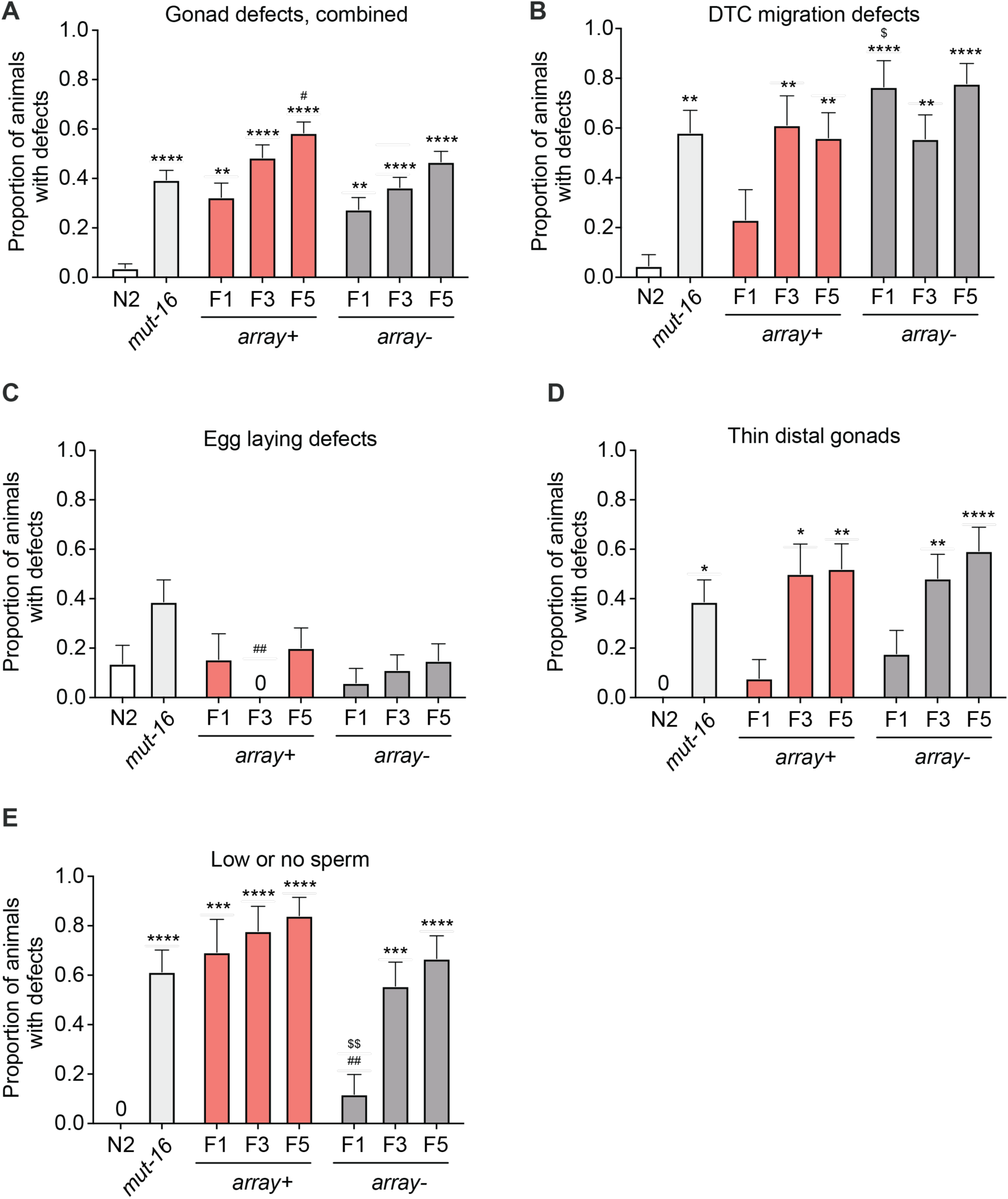
*mut-16* hermaphrodites exhibit numerous gonadal defects. The bar graphs represent mean proportion of animals with germline defects ± S.E.M. in wild type N2, *mut-16(pk710),* and F1, F3, and F5 generations of *mut-16(pk710) array+* and *array-* strains. The germline defects examined were **(A)** total gonadal defects, **(B)** DTC migration defects, **(C)** egg laying defects, **(D)** thin distal gonads, and **(E)** low or no sperm. n ≥ 25 gonad arms over 3 biologically independent trials. *, #, $ *p* < 0.05; **, ##, $$ *p* < 0.01; ***, ###, $$$ *p* < 0.001; ****, ####, $$$$ *p* < 0.0001; one-way ANOVA with Tukey’s *post hoc* test. * denotes a comparison to N2; # denotes a comparison to *mut-16(pk710),* $ denotes a comparison to *array+* within the same generation. Additional data are provided in Table S13.

**Figure S5.**
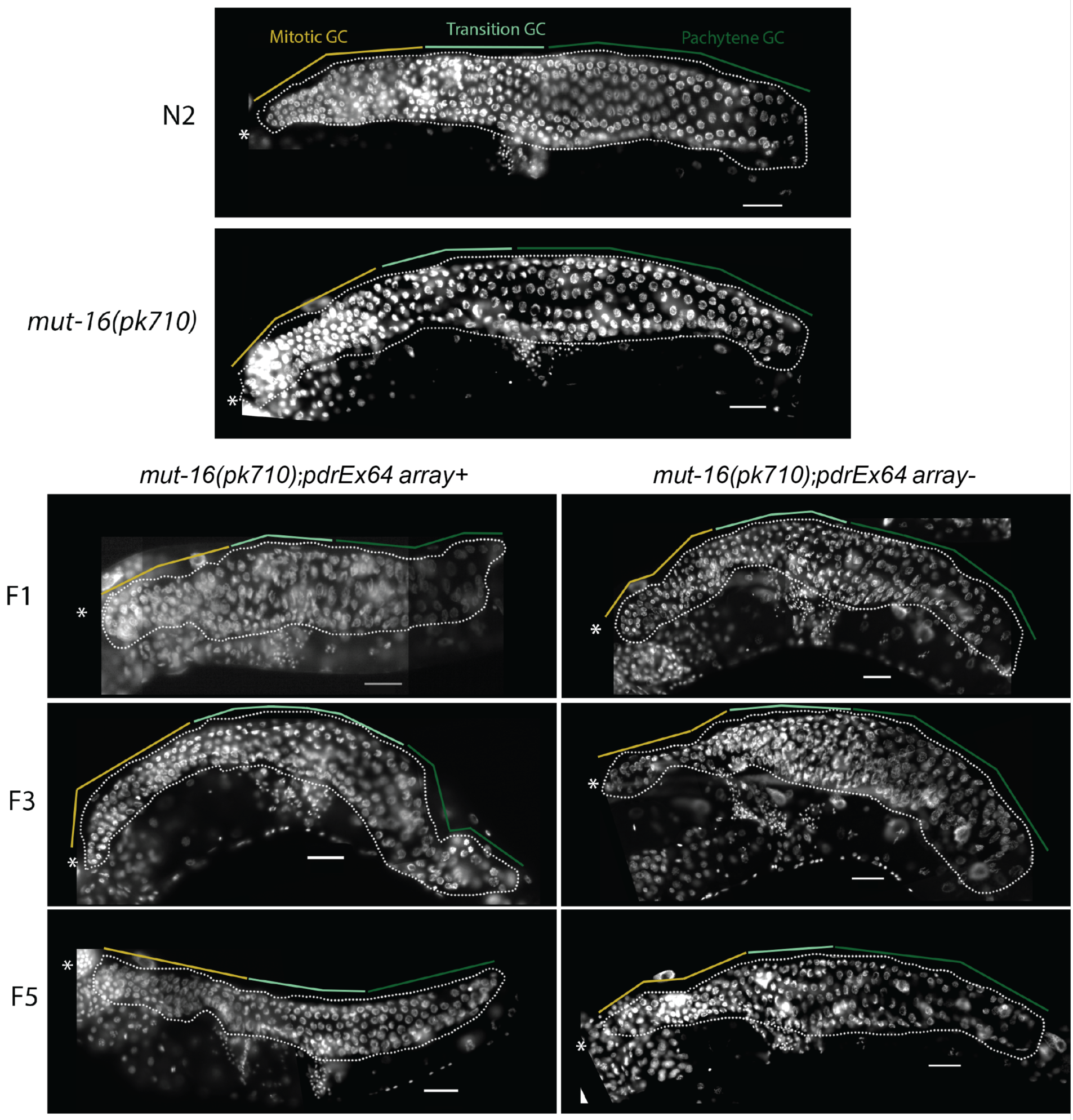
Pan-neuronal overexpression of *mut-16* correlates with decreased germ cell number. Figure shows representative images of DAPI-stained whole mount gonads of wild type N2, *mut-16(pk710),* and F1, F3, and F5 generations of the *mut-16(pk710) array+* and *array-* strains. The yellow, light green, and dark green lines indicate the mitotic, transition, and pachytene germline zones, respectively. The white dotted line demarks the part of the germ line that includes the mitotic, leptotene, zygotene, and pachytene zones of young adult animals. Asterisks indicate the DTC. Scale bar is 25 μm.

**Figure S6.**
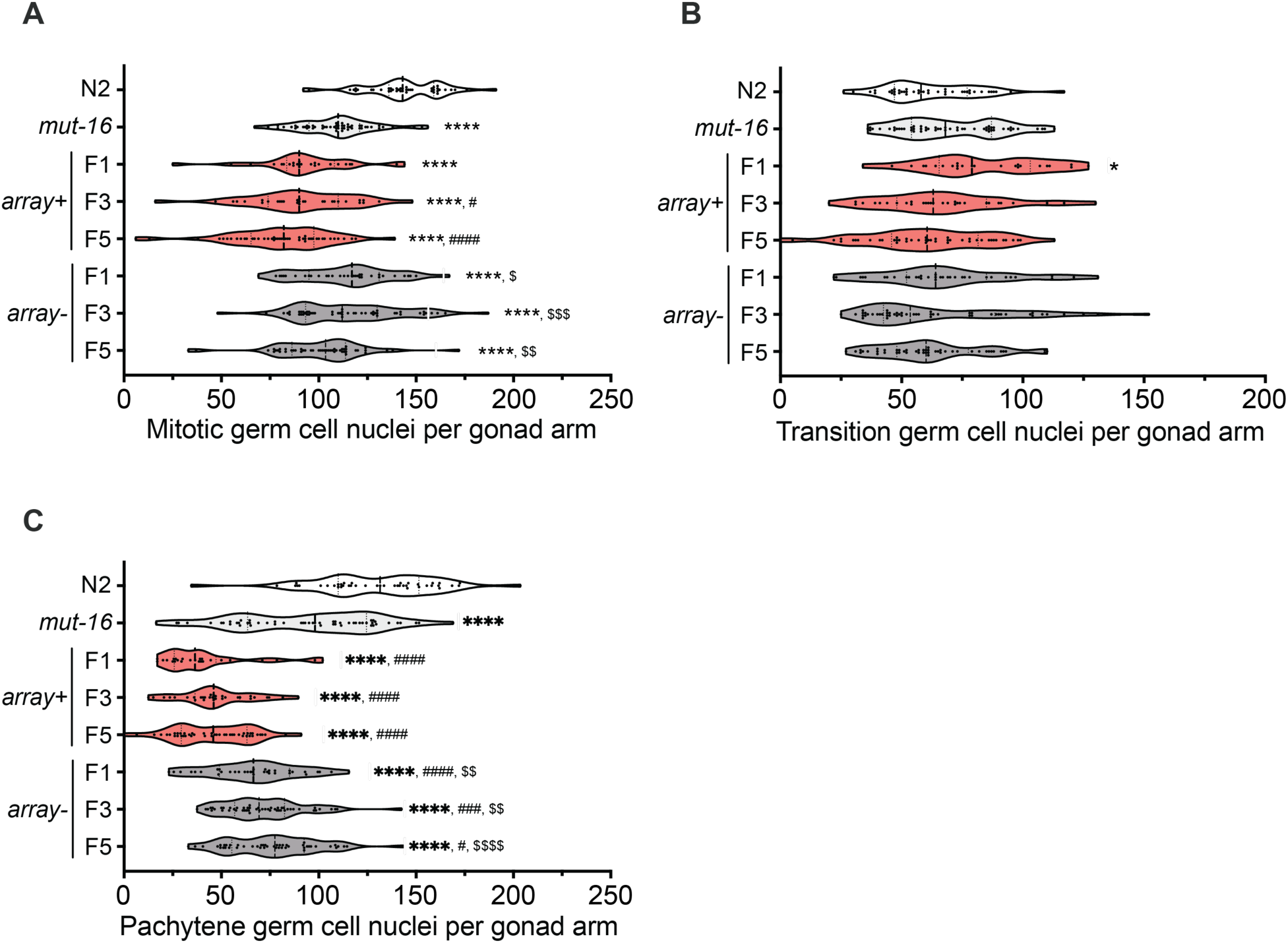
Pan-neuronal overexpression of *mut-16* is correlated with reduced germ cell number in mitotic and pachytene zones. Violin plots represent the number of germ cell nuclei per gonad arm for wild type N2, *mut-16(pk710),* and F1, F3, and F5 generations of *mut-16(pk710) array+* and *array-* strains. The germ cell counts shown are for **(A)** mitotic, **(B)** transition, **(C)** and pachytene zones. n ≥ 25 gonad arms over 3 biologically independent trials. *, #, $ *p* < 0.05; **, ##, $$ *p* < 0.01; ***, ###, $$$ *p* < 0.001; ****, ####, $$$$ *p* < 0.0001; one-way ANOVA with Tukey’s *post hoc* test * denotes a comparison to N2; # denotes a comparison to *mut-16(pk710),* $ denotes a comparison to *array+* within the same generation. Additional data provided in Table S13.

## LITERATURE CITED

Agrawal A. F., 2001 Sexual selection and the maintenance of sexual reproduction. Nature 411: 692–695. https://doi.org/10.1038/35079590

Ailion M, Thomas JH. Dauer formation induced by high temperatures in Caenorhabditis elegans. Genetics. 2000 Nov 1;156(3):1047–67.

Albert P. S., and D. L. Riddle, 1983 Developmental alterations in sensory neuroanatomy of the Caenorhabditis elegans dauer larva. J. Comp. Neurol. 219: 461–481. https://doi.org/10.1002/cne.902190407

Bargmann, C.I. Chemosensation in *C. elegans* (October 25, 2006), WormBook, ed. The C. elegans Research Community, WormBook, doi/10.1895/wormbook.1.123.1, http://www.wormbook.org

Bharadwaj P. S., and S. E. Hall, 2017 Endogenous RNAi Pathways Are Required in Neurons for Dauer Formation in Caenorhabditis elegans. Genetics 205: 1503– 1516. https://doi.org/10.1534/genetics.116.195438

Bhattacharya A., U. Aghayeva, E. G. Berghoff, and O. Hobert, 2019 Plasticity of the Electrical Connectome of C. elegans. Cell 176: 1174–1189.e16. https://doi.org/10.1016/j.cell.2018.12.024

Burt A., 2000 Perspective: Sex, recombination, and the efficacy of selection-Was Weismann right? Evolution (N. Y). 54: 337–351. https://doi.org/10.1111/j.0014-3820.2000.tb00038.x

Butlin R. The costs and benefits of sex: new insights from old asexual lineages. Nature Reviews Genetics. 2002 Apr;3(4):311–7.

Campbell R. J., and H. F. Linskens, 1984 Temperature effects on self incompatibility in Lilium longiflorum. Theor. Appl. Genet. 68: 259–264. https://doi.org/10.1007/BF00266900

Cassada R. C., and R. L. Russell, 1975 The dauerlarva, a post-embryonic developmental variant of the nematode Caenorhabditis elegans. Dev. Biol. 46: 326–342. https://doi.org/10.1016/0012-1606(75)90109-8

Chaudhuri J., V. Kache, and A. Pires-Dasilva, 2011 Regulation of sexual plasticity in a nematode that produces males, females, and hermaphrodites. Curr. Biol. 21: 1548–1551. https://doi.org/10.1016/j.cub.2011.08.009

Devanapally S., S. Ravikumar, and A. M. Jose, 2015 Double-stranded RNA made in C. Elegans neurons can enter the germline and cause transgenerational gene silencing. Proc. Natl. Acad. Sci. U. S. A. 112: 2133–2138. https://doi.org/10.1073/pnas.1423333112

Evans, T. C., ed. Transformation and microinjection (April 6, 2006), WormBook, ed. The C. elegans Research Community, WormBook, doi/10.1895/wormbook.1.108.1, http://www.wormbook.org

Fagan K. A., J. Luo, R. C. Lagoy, F. C. Schroeder, D. R. Albrecht, et al., 2018 A Single-Neuron Chemosensory Switch Determines the Valence of a Sexually Dimorphic Sensory Behavior. Curr. Biol. 28: 902–914.e5. https://doi.org/10.1016/j.cub.2018.02.029

Félix M. A., 2004 Alternative morphs and plasticity of vulval development in a rhabditid nematode species. Dev. Genes Evol. 214: 55–63. https://doi.org/10.1007/s00427-003-0376-y

Francis R., M. K. Barton, J. Kimble, and T. Schedl, 1995 gld-1, a tumor suppressor gene required for oocyte development in Caenorhabditis elegans. Genetics 139.

Frézal L, Félix MA. The natural history of model organisms: C. elegans outside the Petri dish. Elife. 2015 Mar 30;4:e05849.

Gerisch B, Antebi A. Hormonal signals produced by DAF-9/cytochrome P450 regulate C. elegans dauer diapause in response to environmental cues. Development. 2004 Apr 15;131(8):1765–76.

Gibson AK, Delph LF, Lively CM. The two-fold cost of sex: experimental evidence from a natural system. Evolution letters. 2017 May;1(1):6–15.

Golden J. W., and D. L. Riddle, 1982 A pheromone influences larval development in the nematode Caenorhabditis elegans. Science 218: 578–80. https://doi.org/10.1126/SCIENCE.6896933

Gray J. M., J. J. Hill, and C. I. Bargmann, 2005 A circuit for navigation in Caenorhabditis elegans. Proc. Natl. Acad. Sci. U. S. A. 102: 3184–3191. https://doi.org/10.1073/pnas.0409009101

Hall S. E., M. Beverly, C. Russ, C. Nusbaum, and P. Sengupta, 2010 A Cellular Memory of Developmental History Generates Phenotypic Diversity in C. elegans. Curr. Biol. 20: 149–155. https://doi.org/10.1016/j.cub.2009.11.035

Hilliard M. A., C. I. Bargmann, and P. Bazzicalupo, 2002 C. elegans responds to chemical repellents by integrating sensory inputs from the head and the tail. Curr. Biol. 12: 730–734. https://doi.org/10.1016/S0960-9822(02)00813-8

Hodgkin J., H. R. Horvitz, and S. Brenner, 1979 Nondisjunction Mutants of the Nematode Caenorhabditis elegans. Genetics 91: 67–94.

Hussey R, Stieglitz J, Mesgarzadeh J, Locke TT, Zhang YK, Schroeder FC, Srinivasan S. Pheromone-sensing neurons regulate peripheral lipid metabolism in Caenorhabditis elegans. PLoS genetics. 2017 May 18;13(5):e1006806.

Jang H., K. Kim, S. J. Neal, E. Macosko, D. Kim, et al., 2012 Neuromodulatory State and Sex Specify Alternative Behaviors through Antagonistic Synaptic Pathways in C. elegans. Neuron 75: 585–592. https://doi.org/10.1016/j.neuron.2012.06.034

Johnson A. G., 1971 Factors affecting the degree of self-incompatibility in inbred lines of Brussels sprouts. Euphytica 20: 561–573. https://doi.org/10.1007/BF00034212

Kaplan F., J. Srinivasan, P. Mahanti, R. Ajredini, O. Durak, et al., 2011 Ascaroside expression in Caenorhabditis elegans is strongly dependent on diet and developmental stage. PLoS One 6. https://doi.org/10.1371/journal.pone.0017804

Karp X., 2016 Working with dauer larvae *. https://doi.org/10.1895/wormbook.1.180.1

Ketting R. F., T. H. A. Haverkamp, H. G. A. M. Van Luenen, and R. H. A. Plasterk, 1999 mut-7 of C. elegans, required for transposon silencing and RNA interference, is a homolog of werner syndrome helicase and RNaseD. Cell 99: 133–141. https://doi.org/10.1016/S0092-8674(00)81645-1

LeBlanc G. A., and E. K. Medlock, 2015 Males on demand: the environmental-neuro-endocrine control of male sex determination in daphnids. FEBS J. 282: 4080–4093. https://doi.org/10.1111/febs.13393

Lehtonen J., M. D. Jennions, and H. Kokko, 2012 The many costs of sex. Trends Ecol. Evol. 27: 172–178.

Levin D. A., 2012 Mating system shifts on the trailing edge. Ann. Bot. 109: 613–620. https://doi.org/10.1093/aob/mcr159

Lipton J., G. Kleemann, R. Ghosh, R. Lints, and S. W. Emmons, 2004 Mate searching in Caenorhabditis elegans: A genetic model for sex drive in a simple invertebrate. J. Neurosci. 24: 7427–7434. https://doi.org/10.1523/JNEUROSCI.1746-04.2004

Ludewig A. H., and F. C. Schroeder, 2013 Ascaroside signaling in C. elegans. WormBook 1–22. https://doi.org/10.1895/wormbook.1.155.1

McKnight K, Hoang HD, Prasain JK, Brown N, Vibbert J, Hollister KA, Moore R, Ragains JR, Reese J, Miller MA. Neurosensory perception of environmental cues modulates sperm motility critical for fertilization. Science. 2014 May 16;344(6185):754–7.

Meirmans S., P. G. Meirmans, and L. R. Kirkendall, 2012 The costs of sex: Facing real-world complexities. Q. Rev. Biol. 87: 19–40.

Morran L. T., B. J. Cappy, J. L. Anderson, and P. C. Phillips, 2009 Sexual partners for the stressed: facultative outcrossing in the self-fertilizing nematode Caenorhabditis elegans. Evolution 63: 1473–82. https://doi.org/10.1111/j.1558-5646.2009.00652.x

Müller CB, Williams IS, Hardie J. 2001. The role of nutrition, crowding and interspecific interactions in the development of winged aphids. Ecol. Entomol. 26, 330–340. (10.1046/j.1365-2311.2001.00321.x)

Oakley CG, Moriuchi KS, Winn AA. The maintenance of outcrossing in predominantly selfing species: ideas and evidence from cleistogamous species. Annu. Rev. Ecol. Evol. Syst.. 2007 Dec 1;38:437–57.

Otto S. P., 2009 The evolutionary enigma of sex. Am. Nat. 174 Suppl: S1–S14. https://doi.org/10.1086/599084

Ow M. C., K. Borziak, A. M. Nichitean, S. Dorus, and S. E. Hall, 2018 Early experiences mediate distinct adult gene expression and reproductive programs in Caenorhabditis elegans. PLoS Genet. 14: e1007219. https://doi.org/10.1371/journal.pgen.1007219

Paland S., and M. Lynch, 2006 Transitions to asexuality result in excess amino acid substitutions. Science (80-.). 311: 990–992. https://doi.org/10.1126/science.1118152

Pekar O, Ow MC, Hui KY, Noyes MB, Hall SE, Hubbard EJ. Linking the environment, DAF-7/TGFβ signaling and LAG-2/DSL ligand expression in the germline stem cell niche. Development. 2017 Aug 15;144(16):2896–906.

Phillips C. M., T. A. Montgomery, P. C. Breen, and G. Ruvkun, 2012 MUT-16 promotes formation of perinuclear Mutator foci required for RNA silencing in the C. elegans germline. Genes Dev. 26: 1433–1444. https://doi.org/10.1101/gad.193904.112

Pradhan S., S. Quilez, K. Homer, and M. Hendricks, 2019 Environmental Programming of Adult Foraging Behavior in C. elegans. Curr. Biol. 29: 2867–2879.e4. https://doi.org/10.1016/j.cub.2019.07.045

Procko C., Y. Lu, and S. Shaham, 2011 Glia delimit shape changes of sensory neuron receptive endings in C. elegans. Development 138: 1371–1381. https://doi.org/10.1242/dev.058305

Qiao L., J. L. Lissemore, P. Shu, A. Smardon, M. B. Gelber, et al., 1995 Enhancers of Glp-1, a Gene Required for Cell-Signaling in Caenorhabditis elegans, Define a Set of Genes Required for Germline Development. Genetics Society of America.

Rogers A. K., and C. M. Phillips, 2020 RNAi pathways repress reprogramming of C. elegans germ cells during heat stress. Nucleic Acids Res. 48: 4256–4273. https://doi.org/10.1093/nar/gkaa174

Schroeder N. E., R. J. Androwski, A. Rashid, H. Lee, J. Lee, et al., 2013 Dauer-specific dendrite arborization in C. elegans is regulated by KPC-1/furin. Curr. Biol. 23: 1527–1535. https://doi.org/10.1016/j.cub.2013.06.058

Shakes D. C., J. C. Wu, P. L. Sadler, K. LaPrade, L. L. Moore, et al., 2009 Spermatogenesis-specific features of the meiotic program in Caenorhabditis elegans. PLoS Genet. 5: 1000611. https://doi.org/10.1371/journal.pgen.1000611

Sijen T., and R. H. A. Plasterk, 2003 Transposon silencing in the Caenorhabditis elegans germ line by natural RNAi. Nature 426: 310–314. https://doi.org/10.1038/nature02107

Siller S., 2001 Sexual selection and the maintenance of sex. Nature 411: 689–692. https://doi.org/10.1038/35079578

Simon J. M., and P. W. Sternberg, 2002 Evidence of a mate-finding cue in the hermaphrodite nematode Caenorhabditis elegans. Proc. Natl. Acad. Sci. U. S. A. 99: 1598–1603.

Sims J. R., M. C. Ow, M. A. Nishiguchi, K. Kim, P. Sengupta, et al., 2016 Developmental programming modulates olfactory behavior in C. elegans via endogenous RNAi pathways. Elife 5. https://doi.org/10.7554/eLife.11642

Smith J. M., 1971 What use is sex? Journal of Theoretical Biology 30: 319–335. https://doi.org/10.1016/0022-5193(71)90058-0

Smith J. M., 1978 The evolution of sex. (Vol. 4) Cambridge: Cambridge University Press.

Soto M. C., H. Qadota, K. Kasuya, M. Inoue, D. Tsuboi, et al., 2002 The GEX-2 and GEX-3 proteins are required for tissue morphogenesis and cell migranons in C. elegans. Genes Dev. 16: 620–632. https://doi.org/10.1101/gad.955702

Srinivasan J., F. Kaplan, R. Ajredini, C. Zachariah, H. T. Alborn, et al., 2008 A blend of small molecules regulates both mating and development in Caenorhabditis elegans. Nature 454: 1115–1118. https://doi.org/10.1038/nature07168

Tabara H., M. Sarkissian, W. G. Kelly, J. Fleenor, A. Grishok, et al., 1999 The rde-1 gene, RNA interference, and transposon silencing in C. elegans. Cell 99: 123–132. https://doi.org/10.1016/S0092-8674(00)81644-X

Teotónio H., D. Manoel, and P. C. Phillips, 2006 GENETIC VARIATION FOR OUTCROSSING AMONG CAENORHABDITIS ELEGANS ISOLATES. Evolution (N. Y). 60: 1300–1305. https://doi.org/10.1111/j.0014-3820.2006.tb01207.x

Thomas JH, Birnby DA, Vowels JJ. Evidence for parallel processing of sensory information controlling dauer formation in Caenorhabditis elegans. Genetics. 1993 Aug 1;134(4):1105–17.

Uebel C. J., D. C. Anderson, L. M. Mandarino, K. I. Manage, S. Aynaszyan, et al., 2018 Distinct regions of the intrinsically disordered protein MUT-16 mediate assembly of a small RNA amplification complex and promote phase separation of Mutator foci. PLOS Genet. 14: e1007542. https://doi.org/10.1371/journal.pgen.1007542

Webster A. K., J. M. Jordan, J. D. Hibshman, R. Chitrakar, and L. Ryan Baugh, 2018 Transgenerational effects of extended dauer diapause on starvation survival and gene expression plasticity in Caenorhabditis elegans. Genetics 210: 263–274. https://doi.org/10.1534/genetics.118.301250

Weismann A., 1889 The significance of sexual reproduction in the theory of natural selection. E. B. Poulton, S. Schönland, A. E. Shipley, eds. Essays upon Hered. kindred Biol. Probl. Clarendon Press. Oxford. 251–332.

Wexler L. R., R. M. Miller, and D. S. Portman, 2020 C. elegans Males Integrate Food Signals and Biological Sex to Modulate State-Dependent Chemosensation and Behavioral Prioritization. Curr. Biol. 30: 2695–2706.e4. https://doi.org/10.1016/j.cub.2020.05.006

Whitlock M. C., 2000 Fixation of new alleles and the extinction of small populations: drift load, beneficial alleles, and sexual selection. Evolution (N. Y). 54: 1855–1861. https://doi.org/10.1111/j.0014-3820.2000.tb01232.x

Whitlock M. C., and A. F. Agrawal, 2009 Purging the genome with sexual selection: Reducing mutation load through selection on males. Evolution (N. Y). 63: 569–582. https://doi.org/10.1111/j.1558-5646.2008.00558.x

Wong M. C., and J. E. Schwarzbauer, 2012 Gonad morphogenesis and distal tip cell migration in the Caenorhabditis elegans hermaphrodite. Wiley Interdiscip. Rev. Dev. Biol. 1: 519–531.

Yemini E, Jucikas T, Grundy LJ, Brown AE, Schafer WR. A database of Caenorhabditis elegans behavioral phenotypes. Nature methods. 2013 Sep;10(9):877–9.

Zhang C., T. A. Montgomery, H. W. Gabel, S. E. J. Fischer, C. M. Phillips, et al., 2011 mut-16 and other mutator class genes modulate 22G and 26G siRNA pathways in Caenorhabditis elegans. Proc. Natl. Acad. Sci. U. S. A. 108: 1201–1208. https://doi.org/10.1073/pnas.1018695108

